# Impact of the COVID-19 Pandemic on Undergraduate Research in the Department of Biology at Western University: Effect on project types, learning outcomes, and student perceptions

**DOI:** 10.1101/2024.01.24.577125

**Authors:** Ava Chaplin, Susanne Kohalmi, Anne Simon

**Affiliations:** Department of Pathology and Laboratory Medicine, Schulich School of Medicine and Dentistry, Western University, London Ontario; Department of Biology, Faculty of Science, Western University, London Ontario

**Keywords:** undergraduate research, science education, One Health, COVID-19 pandemic, stakeholders, biology

## Abstract

Undergraduate research is a high impact practice that offers numerous benefits to students, academic institutions, and the wider scientific community. Unfortunately, undergraduate research has faced restrictions due to the COVID-19 pandemic. This study aimed to assess how the COVID-19 pandemic has impacted: (1) the number and types of undergraduate research projects performed in the Department of Biology at the University of Western Ontario, and (2) the satisfaction-levels and perceived learning outcomes of students performing these projects. This study also aimed to incorporate a ‘One Health’ framework through an emphasis on stakeholder involvement and the need for future action.

A survey of 33 students who completed an undergraduate research project in the Department of Biology in the 2020/2021 academic year, and 68 students who completed an undergraduate research project in the 5 years prior was conducted. In keeping with the One Health approach, key stakeholders were identified, and a stakeholder map was constructed.

The number of projects performed did not change dramatically despite COVID-19 restrictions. However, a shift towards dry research was observed with 87.9% of students in the 2020/2021 academic year conducting dry research, compared to 16.4% of students in the 5 years prior. Students who conducted research in the 2020/2021 academic year indicated lower overall levels of satisfaction and enjoyment, though their perceived learning outcomes were consistent with students who completed their projects in the 5 years prior. 53 key stakeholders from academia, government, industry, media, and the public were identified.

Students provided invaluable feedback on their undergraduate research experiences that can be used to improve the quality of undergraduate research courses in the Department of Biology in the future. Findings may be of use to other departments and educational institutions that are seeking to improve their own undergraduate research courses amidst the COVID-19 pandemic or looking to incorporate experiential-based learning techniques into existing online courses.

## 1. Introduction

Oftentimes an integral component of the science undergraduate experience is the completion of an independent research project in third or fourth year. These independent research projects typically involve working with a faculty member who acts as a supervisor to the student; performing experimental research; and submitting a detailed written report and/or oral presentation at the end of the course. Undergraduate research plays a pivotal role in allowing students to apply theoretical knowledge to practical skills, and gain experience valued by employers (Deveau, Wang, & Small, 2020; Linn et al., 2015; Lopatto, 2007; Lopatto & Tobias, 2010; Russell, Hancock, & McCullough, 2007; Seymour et al., 2009; Murray, 2018). Undergraduate research also benefits academic institutions by leading to increased publications and presentations, and by enhancing visibility of the scientific community.

The onset of the COVID-19 pandemic resulted in closures of post-secondary academic institutions across the world. Following these closures, many institutions adopted online or blended models of education in order to safely deliver course content and adhere to local, regional, or national COVID-19 guidelines. This unfortunately has meant that undergraduate research courses have faced heavy restrictions, with many students required to conduct their research remotely or having reduced access to on-campus laboratories and research facilities (Deveau, Wang, & Small, 2020; Korbel & Stegle, 2020; Radecki & Schonfeld, 2020; Speer, Lyon, & Johnson, 2021; Trego, Nadybal, Morales, Collins, & Grineski, 2020; Wang, Bauer, Burmeister, Hanauer, & Graham, 2020; Zewail-Foote, 2020). This has also challenged course and program administrators to come up with creative solutions to offer students high quality research-based learning opportunities, despite COVID-19 restrictions.

Prior to the 2020/2021 academic year, the Department of Biology at the University of Western Ontario offered two undergraduate research courses: (1) Bio4999E: Honors Research Thesis, a 1.5 credit research course that spanned the entire academic year (September-April), or (2) Bio4970F/G: Independent Study in Biology, a 0.5 credit independent study course that could be taken in either the fall (September-December) or winter (January-April) semester.

Typically 60-70 students enrolled in Bio4999E per year. While planning for the 2020/2021 academic year, the Department of Biology was concerned that they would not be able to accommodate this number of students as laboratory access on campus had been heavily reduced as a result of the COVID-19 pandemic. Furthermore, the Department of Biology anticipated a second wave of COVID-19 in early-to mid-winter that could cause additional laboratory closures and disruptions to research. Thus, to accommodate students who were still interested in completing a research project over the full academic year despite the COVID-19 pandemic, the Department of Biology at Western University chose to offer students two options:

1. Students could choose to enroll in ‘Bio4999E: Honors Research Thesis’ and receive 1.5 credits. However, if students selected this option they were required to complete their project entirely remotely (either *in silico* or field based).
2. Students could choose to take ‘Bio4970F: Independent Study in Biology’ in the fall semester, followed by a newly developed course, ‘Bio4971G: Independent Study in Biology’, in the winter semester. If students selected this option, they were permitted to complete experimental research in laboratories on campus, provided that their research supervisors were able to accommodate them and COVID-19 social-distancing guidelines were upheld. Additionally, in the case that COVID-19 related disruptions meant that a student’s research could not move forward as planned, they were not required to continue with Bio4971G after the completion of Bio4970F in the fall. Thus, this option allowed both students and their supervisors flexibility.

With this in mind, the overall objective of this study was to assess and describe the impact of the COVID-19 pandemic on undergraduate research in the Department of Biology at Western University through a One Health perspective. To do so, a survey of students who conducted undergraduate research in the Department of Biology in the 2020/2021 academic year and, to provide a measure of comparison, students who conducted undergraduate research in the Department of Biology in the 5 academic years prior, was performed.

## 2. Background

To describe the context in which this study takes place, and to inform development of the survey, an informal literature review was performed. The key themes identified through this literature review are presented below:

### 2.1 Impact of the COVID-19 Pandemic on Post-Secondary Students

The COVID-19 pandemic has led to a widespread shift to online or blended models of learning in post-secondary education. Online learning exacerbates existing challenges and poses new challenges for students (Castro & Tumibay, 2019; Rasheed, Kamsin, & Abdullah, 2020). In the context of the COVID-19 pandemic, many of these challenges are further amplified as students are forced to navigate online learning while also facing a myriad of COVID-19 related concerns (Adedoyin & Soykan, 2020; Christian, McCarty & Brown, 2020; Gillis & Krull, 2020; Mukhtar et al., 2020). Post-secondary students have reported increased distractions and difficulty concentrating, decreased motivation and time management ability, increased anxiety about the future, increased concern about academic performance, uncertainty over course expectations, and technological difficulties or lack of access to appropriate technology, as barriers to online learning during the COVID-19 pandemic (Adedoyin & Soykan, 2020; Gillis & Krull, 2020; Mukhtar et al., 2020).

Furthermore, students have reported numerous COVID-19 related stressors that extend beyond the classroom. These include, while are not limited to, the loss of social networks, concerns related to their own health or the health of friends and loved-ones, loss of job and internship opportunities, and fears about potential long-term impacts of COVID-19 on career prospects (Aucejo, French, Araya, & Zafar, 2020; Kleiman et al., 2020). For instance, a survey of 1500 students at Arizona State University found that as a result of the COVID-19 pandemic, 13% had delayed graduation, 40% had lost a job, internship, or job offer, and 29% expected to earn less at age 35 (Aucejo, French, Araya, & Zafar, 2020). Due to these factors, it has been proposed that post-secondary students may be particularly vulnerable to experiencing negative mental health consequences amidst the COVID-19 pandemic.

An overwhelming number of studies conducted in various countries worldwide have found that the COVID-19 pandemic has led to increased stress, anxiety, insomnia, loneliness, substance-use, depression, and suicidal ideations among post-secondary students (Aucejo, French, Araya, & Zafar, 2020; Cao et al., 2020; Elmer, Mepham, & Stadtfeld, 2020; Kaparounaki et al., 2020; Kleiman, Yeager, Grove, Kellerman, & Kim, 2020; Meda et al., 2021; Son, Hegde, Smith, Wang, & Sasangohar, 2020; Wang et al., 2020; Wieczorek et al., 2021). For instance, in a survey of 2031 students at Texas A&M University, 71.26% indicated that their stress and anxiety levels had increased during the pandemic, and 48.14% demonstrated symptoms in line with moderate-to-severe levels of depression (Wang et al., 2020). Moreover, less than half of the students (43.25%) indicated that they were able to cope adequately with stress related to the COVID-19 situation, and very few students reported using university health services such as counselling (10.34%) or health services outside of the university (4.38%). These findings highlight an urgent need for attention and support to students experiencing increased mental health burdens associated with the COVID-19 pandemic.

### 2.2 Importance of Undergraduate Research

The benefits of undergraduate research are numerous and far reaching (Deveau, Wang, & Small, 2020; Linn et al., 2015; Lopatto, 2007; Lopatto & Tobias, 2010; Russell, Hancock, & McCullough, 2007; Seymour et al., 2009; Murray, 2018). Indeed, undergraduate research is considered a “high impact practice” (HIP) in post-secondary education, a title that is given to educational practices (of which there are 11 in total) that are thought to lead to enhanced student engagement and elevated learning outcomes (Murray, 2018). The benefits of undergraduate research to students include engaging a wide range of learning preferences, providing practical training and transferable skills, promoting active learning and innovation, enabling intellectual independence, fostering creativity, analytical thinking and problem solving, enhancing career preparation and communication skills, and establishing a growth mindset, among others (Deveau, Wang, & Small, 2020; Linn et al., 2015; Lopatto, 2007; Lopatto & Tobias, 2010; Russell, Hancock, & McCullough, 2007; Seymour et al., 2009; Murray, 2018). Finally, beyond benefiting students, undergraduate research also contributes to knowledge generation and exchange, leads to increased publications and conference presentations for academic institutions, and promotes the visibility of the scientific community. Undergraduate research is thus important to consider in the context of the COVID-19 pandemic, an event that has led to severe disruptions to educational and research environments.

### 2.3 Impact of the COVID-19 Pandemic on Undergraduate Research

There is ample evidence of widespread impacts of the COVID-19 pandemic on research activities (Korbel & Stegle, 2020; Radecki & Schonfeld, 2020). At many universities, all non-essential research was halted at the onset of the pandemic (between January-March 2020). A survey of 881 life scientists from Germany, Spain, the UK, Italy, France, Canada, Turkey and the USA found that by April 2020, 57% of scientists had lost some of their research work (Korbel & Stegle, 2020). 25% reported at least 1 to 6 months of work had been lost, with dramatic differences seen between scientists working in wet labs (73%) and dry labs (31%). Many wet lab researchers were faced with difficult decisions regarding how best to care for mice, fish, monkeys, and other research animals, with some opting for euthanization in order to protect the safety of their staff and to preserve the animals’ remaining welfare (Radecki & Schonfeld, 2020). However, some scientists reported that they enjoyed this “break” from the lab as it allowed them to catch up on readings, writing projects, organizing data, and performing back-burned data analysis. Some scientists also reported that the COVID-19 associated shut-downs promoted increased collaboration and data-sharing between colleagues. While many labs began to reopen in the summer of 2020, they are still operating under heavy restrictions to this day. Indeed, many labs are operating in rotational shifts, with all researchers required to wear masks and other PPE and to remain socially distanced from one another in the lab space (Government of Canada, 2020; Radecki & Schonfeld, 2020). Furthermore, many researchers are required to undergo COVID-19 related training, and complete COVID-19 screenings prior to entering the lab.

Due to these restrictions, many laboratories were unable to accommodate undergraduate researchers for the 2020/2021 academic year, or were only able to offer them projects that could be completed remotely. This has had drastic impacts on the delivery of undergraduate research courses and the experiences of students performing undergraduate research (Deveau, Wang, & Small, 2020; Speer, Lyon, & Johnson, 2021; Trego, Nadybal, Morales, Collins, & Grineski, 2020; Wang, Bauer, Burmeister, Hanauer, & Graham, 2020; Zewail-Foote, 2020). For instance, a study of 983 undergraduate students across 16 universities in the United States found that almost half (44.5%) of undergraduate students who planned to do research during the summer semester had their programs cancelled due to COVID-19 (Trego, Nadybal, Morales, Collins, & Grineski, 2020). Students whose projects were cancelled reported concerns that the loss of the opportunity would impact their ability to get into graduate school or get research-related employment after completing undergrad. Students who did conduct research in summer 2020 reported decreased communication with their faculty mentors as a result of the COVID-19 pandemic, and lower levels of satisfaction with their faculty mentors (though overall levels of satisfaction were still high).

Another study examining the impact of COVID-19 on undergraduate research in the Department of Chemistry and Physics at the University of New England (UNE) found that the loss of physical laboratory experimentation prevented students from gaining manual laboratory skills. This in turn limited students’ ability to grasp conceptual knowledge behind specific procedures, and negatively hindered their troubleshooting skills and ability to evaluate data (Deveau, Wang, & Small, 2020). Furthemore, instructors reported difficulty in keeping students focused during synchronous virtual meetings, and found that technical issues prevented students from fully immersing in the undergraduate research experience, subsequently leading to decreased motivation. The transition to online learning was also reported to have dramatically reduced student-to-student interactions, which subsequently lead to a diminished sense of community and camaraderie among students.

A third study found that students required to transition from in-person to online research and instruction amidst the COVID-19 pandemic expressed disappointment for losing the opportunity to carry out their planned laboratory experiments. They reported decreased levels of ‘situational interest’, which refers to a state of heightened motivation characterized by increased attention, effort and affect (Wang, Bauer, Burmeister, Hanauer, & Graham, 2020). Thus, the COVID-19 pandemic led to a wide range of unforeseen changes to undergraduate research. The findings from studies examining these changes may help universities, faculty supervisors, and students plan and prepare for interruptions in the future.

### 2.4 Relevance to One Health

One Health is a science-based, collaborative approach that aims to safeguard and improve human, animal, and environmental health (Centers for Disease Control and Prevention, 2019.; World Health Organization, 2017). As such, research and education are key to One Health as they allow for knowledge generation and exchange (Gibbs, 2014; Lerner & Berg, 2015). Indeed, research is critical to identifying the most effective ways to promote human, animal and environmental health. Without research, we would be unable to effectively prevent, detect, treat, and control various health issues. Moreover, modern research in the fields of science, technology, engineering or math (STEM) almost always involves collaboration between individuals within and across fields of expertise, honouring the One Health emphasis on interdisciplinary collaboration.

This study specifically pertains to undergraduate research in biology, a field that inherently adheres well to the One Health approach due to its natural integration of human, animal, and environmental health. Research in biology covers a broad range of topics across all three pillars of One Health, including: genetics, cell biology, physiology, biochemistry, synthetic biology, ecology, plant morphology, evolution, and conservation. Biological research often uses animals, plants, or fungi to model human pathophysiologies or to gain insight into our natural environments, providing a direct example of a One Health application in science.

Finally, in the context of the COVID-19 pandemic, post-secondary students have experienced drastic changes to their learning and social environments. This has caused drastic repercussions to the health and wellbeing of students, particularly as it relates to mental health. Thus, this study is relevant to One Health in its focus on research in the field of Biology, and in the context of its significant social and human health impacts of the COVID-19 pandemic on students.

## 3. Specific Objectives

**Objective 1:** Assess the impact of the COVID-19 pandemic on the number and types of undergraduate research projects performed, and the use of animal models in undergraduate research within the Department of Biology at Western University.

**Objective 2:** Assess the impact of the COVID-19 pandemic on the satisfaction and enjoyment levels, and the perceived learning outcomes of students conducting undergraduate research within the Department of Biology at Western University.

**Objective 3:** Identify key stakeholders and emphasize the need for action by developing a list of recommendations to improve the quality of undergraduate research within the Department of Biology at Western University in future years.

## 3. Methods

### 3.1 Study Population

To address these objectives, a survey of students who completed Bio4999E or Bio4970/71 in the 2020/2021 academic year, and students who completed Bio4999E in the 5 preceding academic years, e.g. 2015/2016, 2016/2017, 2017/2018, 2018/2019, or 2019/2020, was conducted. In keeping with the standards set by the Human Research Ethics board at Western University, the names and contact information of potential study participants were identified primarily through publicly available sources. Indeed, the names of students who completed Bio4999E in the 5 preceding academic years were obtained by examining the schedules of the annual in-house biology conference at which Bio4999E students are required to present. As this presentation schedule is prepared in March, it was not available in time for identification of the names of students conducting their projects in the 2020/2021 academic year. Thus, the names of students enrolled in Bio4999E or Bio4970/71 in the 2020/2021 academic year were provided directly by the Department of Biology. The contact information of potential study participants was obtained through Google searches or the Western University Student Directory. Study participants were invited to participate in the study via email or Linkedin.

### 3.2 Survey Design

Two separate surveys were developed: one for the ‘current cohort’, i.e. students who completed their projects in the 2020/2021 academic year, and one for the ‘past cohort’, i.e. students who completed their projects in the 2015/2016, 2016/2017, 2017/2018, 2018/2019, or 2019/2020 academic years (Appendix 1). These surveys were identical apart from slight differences in wording and the addition of a couple questions pertaining to the COVID-19 pandemic in the survey designed for the current cohort.

Surveys were divided into three sections. The first section collected background information on participants, including whether they were enrolled in Bio4999E or Bio4970/71 (for the current cohort), what year they completed their project in (for the past cohort), whether they had prior research experience, and whether their study results had been shared or they anticipate sharing their results through a publication and/or a presentation at a conference other than the year-end Department of Biology Conference. Additionally, the first section of the survey also assessed project type and the usage of animal models. The criteria used to assess project type included (1) whether the research performed was wet (involving experimental work taking place in a laboratory), dry (involving *in silico* or computer-based techniques, such as bioinformatics, data analysis or literature reviews), or field based (involving collection of primary data from natural environments); and (2) the home department of the research supervisors (as an indirect measure of the field the research was conducted in). This section of the survey consisted of closed-ended questions in which participants were required to select from a given number of responses to simplify data input and analysis (Rea & Parker, 2014; Totten, Panacek & Price, 1999).

The second section of the survey assessed participants’ overall level of satisfaction and enjoyment with their undergraduate research experience, and the degree to which they believed to have achieved course learning outcomes. A five-point Likert Scale was used in which participants were asked to select their level of agreement with a series of statements. This allowed for the collection of ordinal data that could be statistically analyzed (Rea & Parker, 2014; Totten, Panacek & Price, 1999). Course learning outcomes that were assessed included the following: skill in writing a research proposal, ability to develop a central hypothesis and goal, ability to search, read, and evaluate primary scientific literature, ability to conduct and troubleshoot research, ability to evaluate and analyze data, skill in writing a progress report and final thesis, and ability to defend scientific data, a research approach, and interpretation of data. The third and final section of the survey consisted of two free response, short-answer questions where participants were invited to provide additional comments about their experience and any specific recommendations they have to improve the quality of Bio4999E or Bio4970/71 courses in the future. This portion of the survey allowed for collection of qualitative data (Rea & Parker, 2014; Totten, Panacek & Price, 1999).

Prior to distribution, the survey was submitted to the Western Research Ethic’s Board for approval. The survey was distributed using the secure platform ‘Qualtrics’ which employs encryption technology and restricted access authorizations to protect all data collected.

### 3.3 Data Analysis

Descriptive statistics were compiled to describe participants’ background in research and type of research conducted. Multiple-choice and Likert-scale responses are presented as percent of responses. Furthermore, Likert scale responses were assigned a value from 1 (strongly disagree) to 5 (strongly agree) and an overall satisfaction and learning outcomes score was calculated per participant. The median, maximum and minimum values from the overall satisfaction score and the learning outcomes score were also calculated. Figures were generated using Prism 8 (GraphPad). Qualitative answers to the two free response questions were analyzed using a preliminary visual inspection approach that involved one individual (AC) sorting through the data and identifying recurrent themes. For future work, qualitative data will be analyzed with thematic analysis, although due to time constraints this was not feasible for the present work (Clarke & Braun, 2016).

### 3.4 Stakeholder Analysis

Stakeholders were identified using Google searches. Various Western University, governmental, research organization, social media, and news outlet websites were searched to compile a list of stakeholders across academia, government, industry, media and the public.

## 4. Results

### 4.1 Participants

Of a total study population of 375 individuals, the contact information of 276 (73.6%) was identified. Of the 276 contacted, 99 (35.9%) responded, representing 26.4% of the total study population. Survey respondents include 33 students who conducted research in the 2020/2021 academic year (current cohort), and 66 students who conducted research in the 5 academic years prior (past cohort) (Table 1). Response rates were higher among the current cohort (33 out of 61 responded, or 54.1%), compared to the past cohort (66 out of 215 responded, or 30.7%). The majority of participants were enrolled in the Biology (44.4%) or Genetics (43.4%) module offered by the Department of Biology, while the remaining were in Genetics and Biochemistry (5.1%), Animal Behaviour (3.0%), Biodiversity and Conservation (2.0%) and Integrated Sciences (1.0%) (Tables 1 & 2).

**Table 1.** Participants’ demographic characteristics.

| Variables | Total Participants (N=99) |
| --- | --- |
| **Academic Year in Which Project Was Completed, n (%)** |  |
|  | 11 (11.1) |
| 2015/2016 | 16 (16.2) |
| 2016/2017 | 10 (10.1) |
| 2017/2018 | 13 (13.1) |
| 2018/2019 | 16 (16.2) |
| 2019/2020 | 33 (33.3) |
| 2020/2021 |  |
| **Module, n (%)** | 44 (44.4) |
| Biology | 43 (43.4) |
| Genetics | 6 (6.1) |
| Genetics and Biochemistry | 3 (3.0) |
| Animal Behaviour | 2 (2.0) |
| Biodiversity and Conservation | 1 (1.0) |
| Integrated Sciences |  |

**Table 2:**
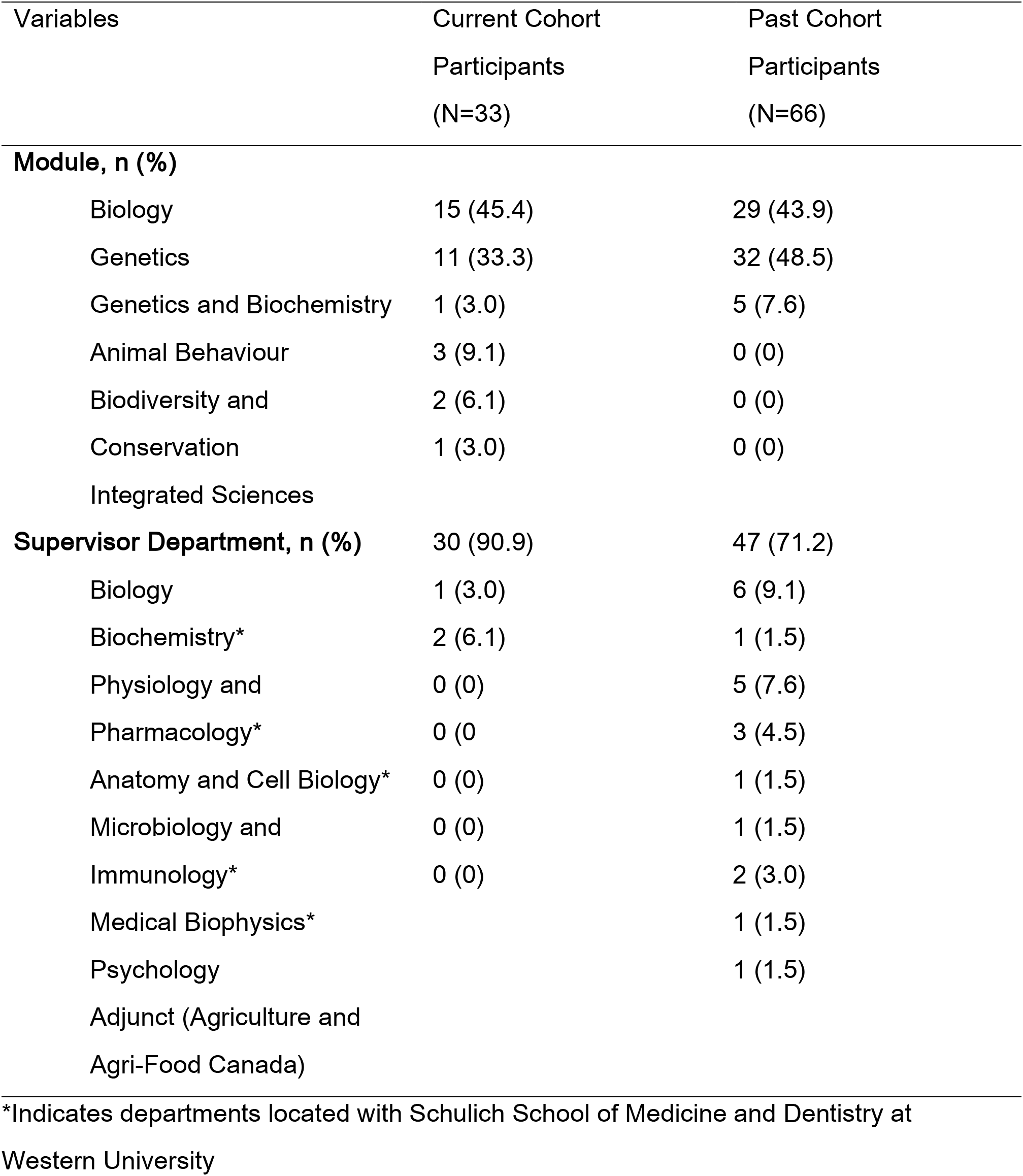
Student modules and research supervisor departments divided by current cohort (2020/2021 academic year) and past cohort (5 academic years prior to 2020/2021)

### 4.2 Number and Types of Projects, and Usage of Animal Models

The number of projects performed did not change dramatically despite COVID-19 related restrictions (Figure 1). A shift towards dry research was observed in students who conducted their projects in the 2020/2021 academic year: 87.8% of students in the current cohort completed dry research, compared to only 15.2% in the past cohort (Figure 2). Despite the shift to dry research in the 2020/2021 academic year, the percentage of students using animal models did not decrease, and in fact a slight increase occurred (Table 3 & Figure 3). 45.5% of the current cohort used animal models, compared to 37.9% of the past cohort. In accordance with the shift to dry research, the majority of projects using animal models in the current cohort were dry (73.3%), whereas the majority of projects using animal models in the past cohort were wet (77.3%). A wide range of animal models were used, including fruit flies, mice, salmonids, butterflies, African impalas, spiders, termites, mites, birds, and beetles (Table 3).

**Figure 1.**
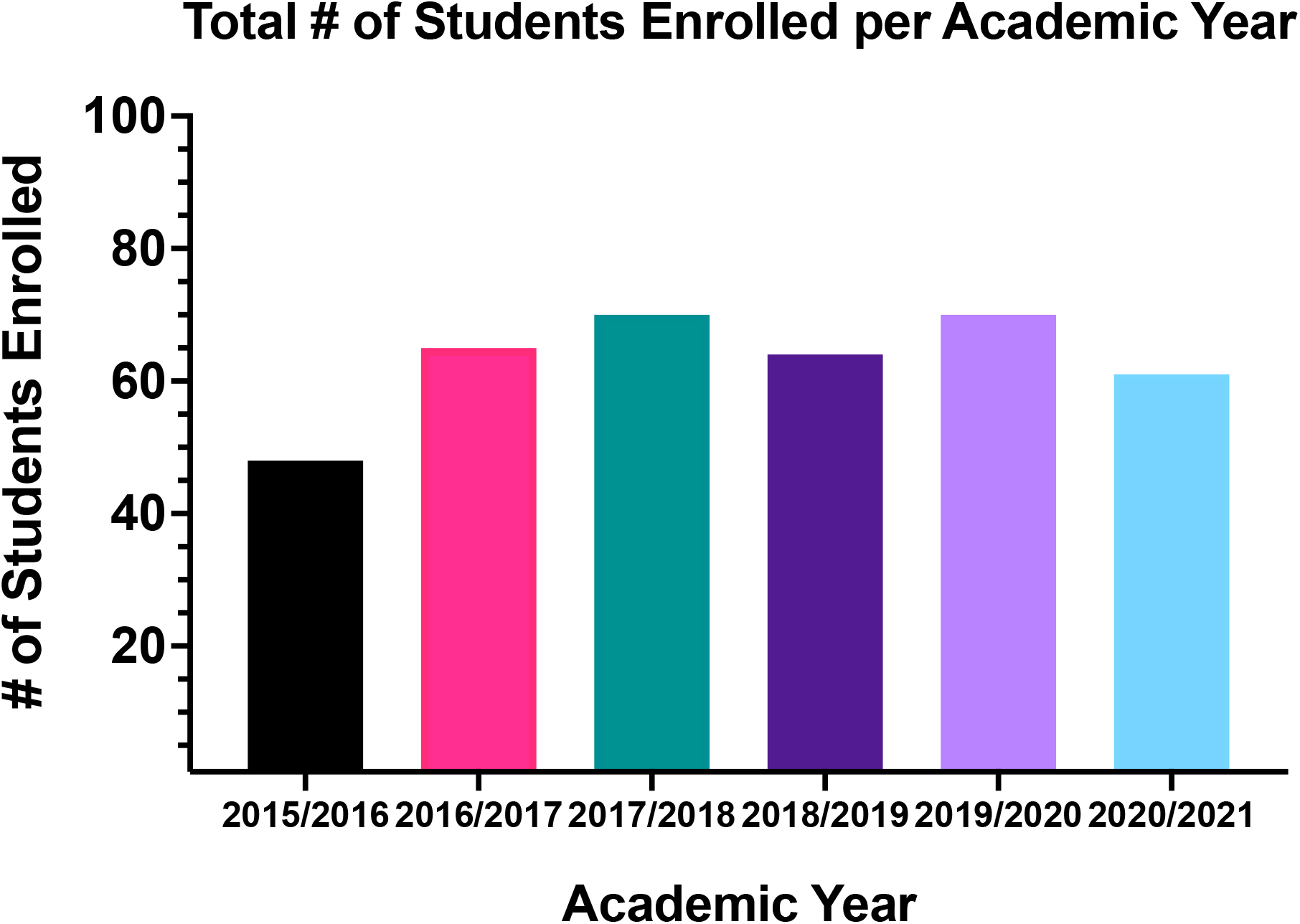
Enrollment in full-year undergraduate research in the Department of Biology per academic year.

**Figure 2.**
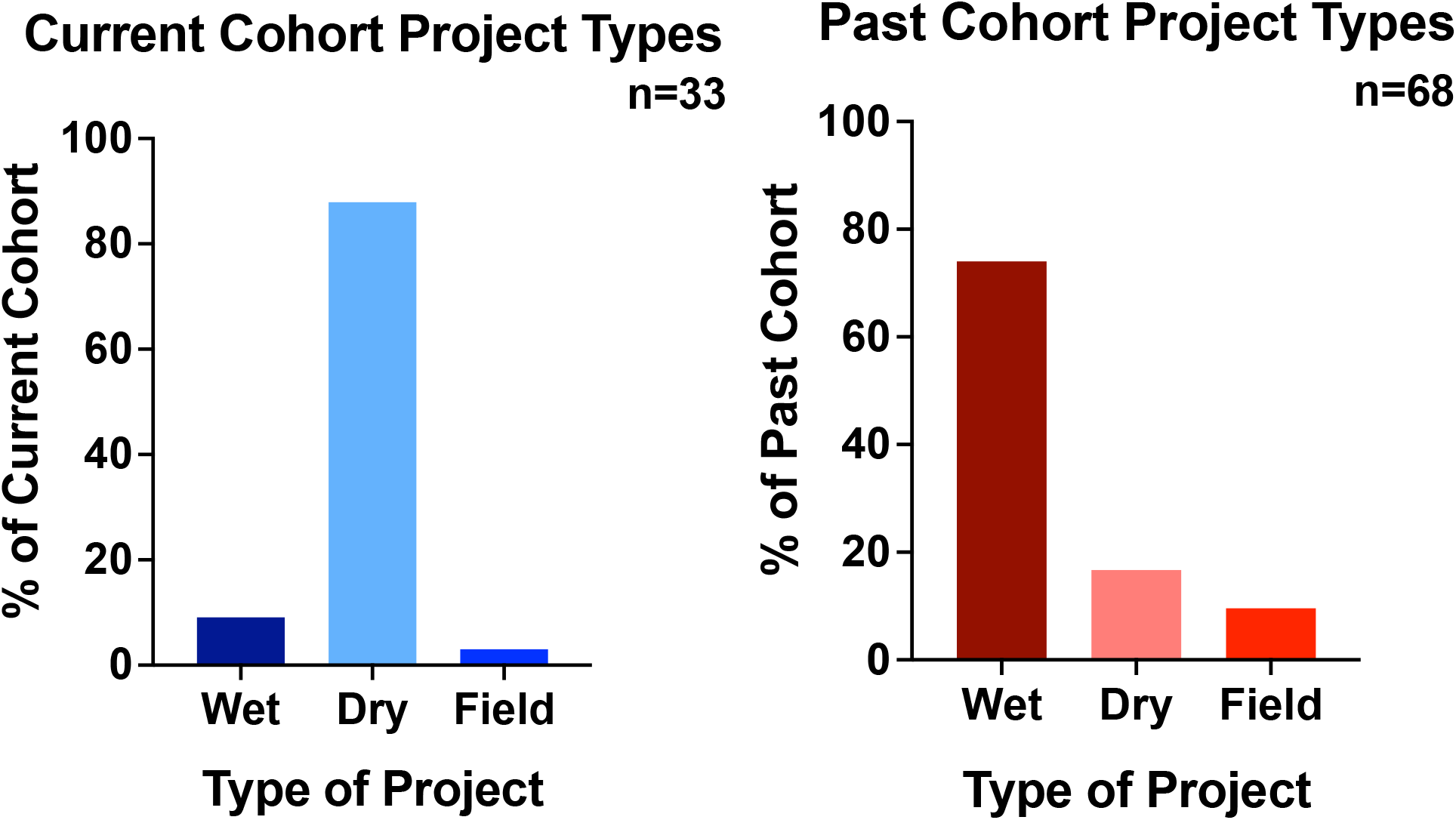
Type of project performed divided by current cohort (2020/2021 academic year) and past cohort (5 academic years prior to 2020/2021)

**Figure 3.**
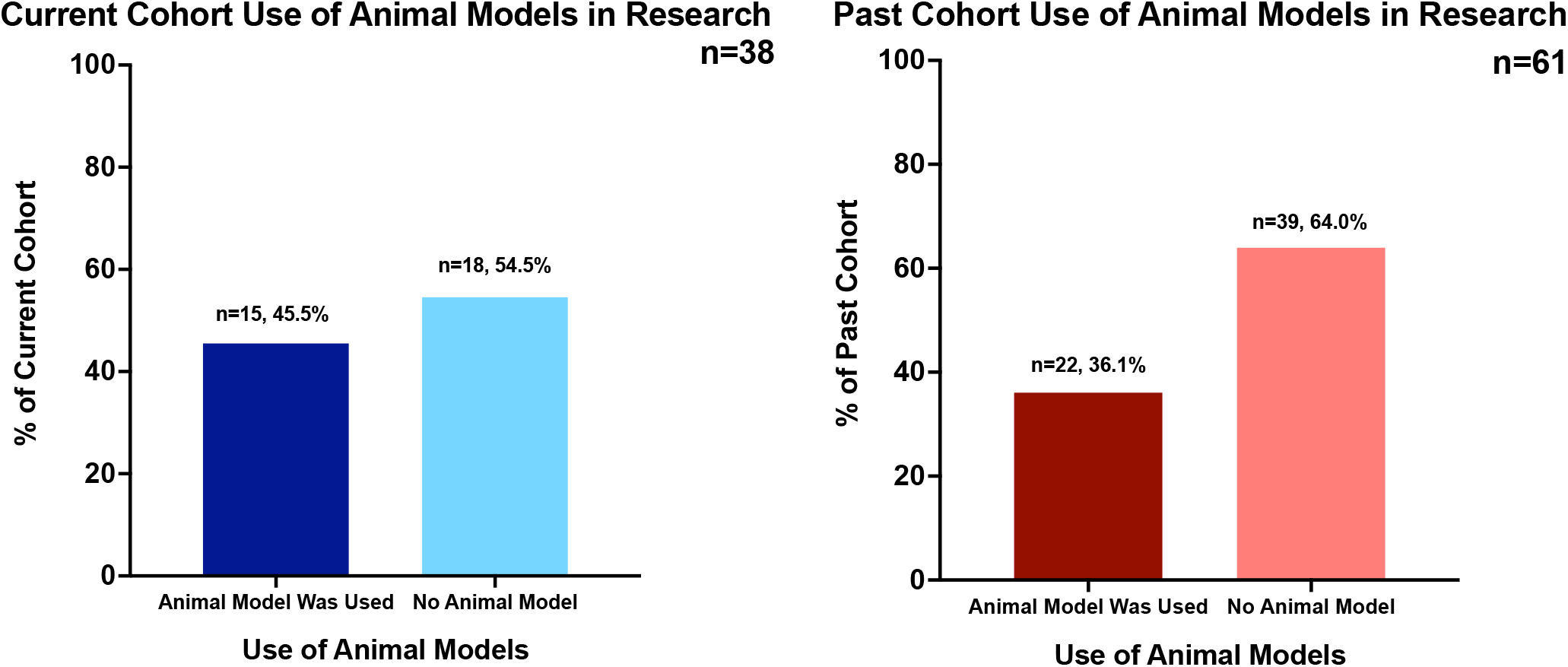
Percentage of projects using animal models divided by current cohort (2020/2021 academic year) and past cohort (5 academic years prior to 2020/2021).

**Table 3:** Usage of animal models divided by current cohort (2020/2021 academic year) and past cohort (5 academic years prior to 2020/2021)

| Animal models used by the<br>current cohort (N=33) | Participant<br>s, n (%) | Animal models used by the<br>past cohort (N=66) | Participants,<br>n (%) |
| --- | --- | --- | --- |
| Eastern subterranean<br>termites | 1 (3.0)<br>2 (6.1) | Easter subterranean<br>termites | 1 (1.5)<br>5 (7.6) |
| Mice | 1 (3.0) | Mice | 2 (1.5) |
| Salmonids | 1 (3.0) | Salmonids | 1 (1.5) |
| Butterflies | 1 (3.0) | Butterflies | 10 (15.1) |
| Black widow spider | 1 (3.0) | Vinegar fly | 1 (1.5) |
| European garden spider | 1 (3.0) | Colorado potato beetle | 2 (1.5) |
| Cellar spider | 1 (3.0) | Ground squirrels | 2 (3.0) |
| Two spotted spider mite | 1 (3.0) | Brown headed cowbirds | 1 (1.5) |
| Crustacean zooplankton | 2 (6.1) | Soil-dwelling mite |  |
| African Impalas | 3 (9.1) |  |  |
| Other (not stated) |  |  |  |
| Total | 15 (45.5) |  | 24 (37.9) |
| : |  |  |  |

90.9% of the current cohort had research supervisors in the Department of Biology, compared to 71.2% of the past cohort (Table 2). Indeed, in the current cohort only 3 students (9.1%) indicated that they had supervisors outside of the Department of Biology, with 2 indicating that their supervisor was in Physiology and Pharmacology at Schulich School of Medicine and Dentistry, and 1 indicating that their supervisor was in the Biochemistry at Schulich. In contrast, in the past cohort 18 students (28.8%) indicated that they had supervisors outside of the Department of Biology, mainly in various departments at Schulich (6 in Biochemistry, 5 in Anatomy and Cell Biology, 3 in Microbiology and Immunology, 1 in Physiology and Pharmacology, and 1 in Medical Biophysics), though 1 student indicated that their supervisor was in the Department of Psychology, and 2 indicated that their supervisors were adjunct professors who worked at Agriculture and Agri-Food Canada.

### 4.3 Satisfaction Levels and Learning Outcomes

Students from the current cohort indicated overall lower levels of satisfaction and enjoyment compared to the past cohort. Indeed, overall 57.6% of the current cohort selected either “Strongly Agree” (20.7%) or “Agree” (36.9%) when asked to rank their agreement to a range of statements pertaining to overall satisfaction and enjoyment (Figure 4). In comparison 87.7% of the past cohort selected either “Strongly Agree” (53.4%) or “Agree” (36.1%) when asked to rank their level of agreement to the same statements. A particularly large drop occurred in the percentage of students that selected “Strongly Agree” (53.4% of the past cohort, compared to 20.7% of the current cohort). The overall satisfaction scores (which could be anywhere between 5-30, with 5 indicating strong dissatisfaction and 30 indicating strong satisfaction) had a median value of 21 for the current cohort, and ranged between a minimum value of 10 to a maximum value of 30. The overall satisfaction scores for the past cohort had a median value of 27, and ranged between 6 to 30.

**Figure 4.**
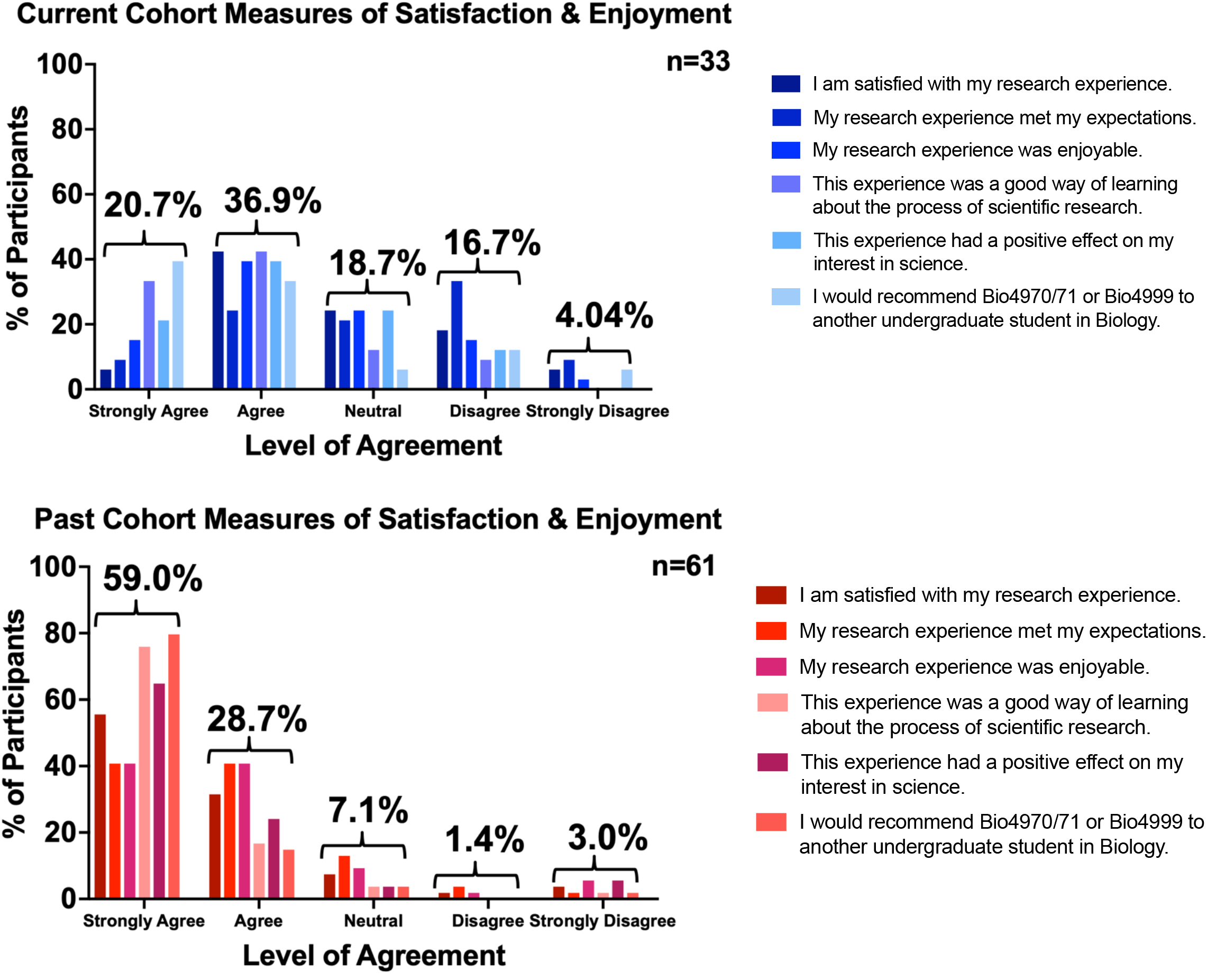
Measures of satisfaction and enjoyment divided by current cohort (2020/2021 academic year) and past cohort (5 academic years prior to 2020/2021)

In section 1 of the current cohort survey, students were asked to state whether their research project underwent any changes between registration in Bio4999 or Bio4970 in June 2020 and the first proposal deadline in October 2020, and were given the option of selecting between major, minor or no project alterations. Examples of major vs. minor changes were provided for clarity (e.g. major alterations included a shift in research focus or a switch from wet to dry research, while minor alterations included an altered hypothesis or updated project goals). Within the current cohort, students who indicated that their project had undergone major alterations reported higher levels of dissatisfaction compared to those who indicated that their project underwent minor or no alterations (Figure 5). Indeed 44.1% of those whose project underwent major alterations selected either “Strongly Agree” (17.9%) or “Agree” (26.2%), while in comparison 71.3% of those whose projects underwent minor or no alterations selected “Strongly Agree” (24.1%) or “Agree” (47.2%). Moreover, the median satisfaction score for those whose projects underwent major alterations was 19.5, compared to 23 for those whose projects underwent minor or no alterations.

**Figure 5.**
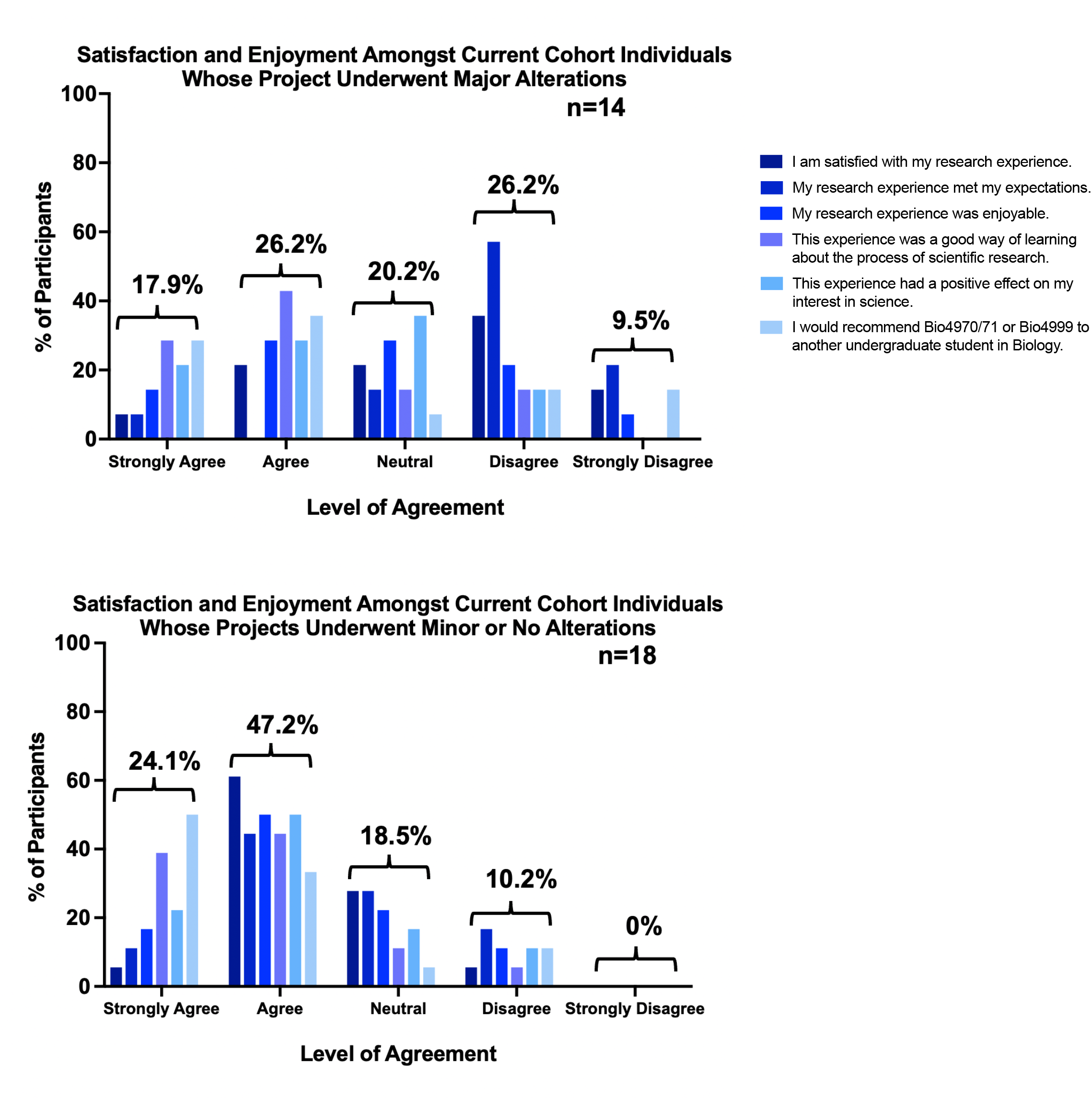
Measures of satisfaction in the current cohort (2020/2021 academic year) between individuals whose project underwent major alterations compared to individuals whose projects underwent minor or no alterations.

Achievement of course learning outcomes remained fairly consistent between the current and past cohorts, with both groups indicating achieving course learning outcomes at high rates (Figure 6). Indeed, 76.2% of the current cohort selected either “Strongly Agree” (36.4%) or “Agree” (39.8%) when asked to rank their level of agreement with a range of statements pertaining to achieving course learning outcomes. In comparison, 89.5% of the past cohort selected either “Strongly Agree” (53.4%) or “Agree” (36.1%). The overall learning outcome score (which could be anywhere between 5-35, with 5 indicating low levels of achievement and 30 indicating high levels) had a median value of 28 for the current cohort and ranged between 19 to 35. The past cohort had a median score of 31, with a minimum of 7 and a maximum of 35.

**Figure 6.**
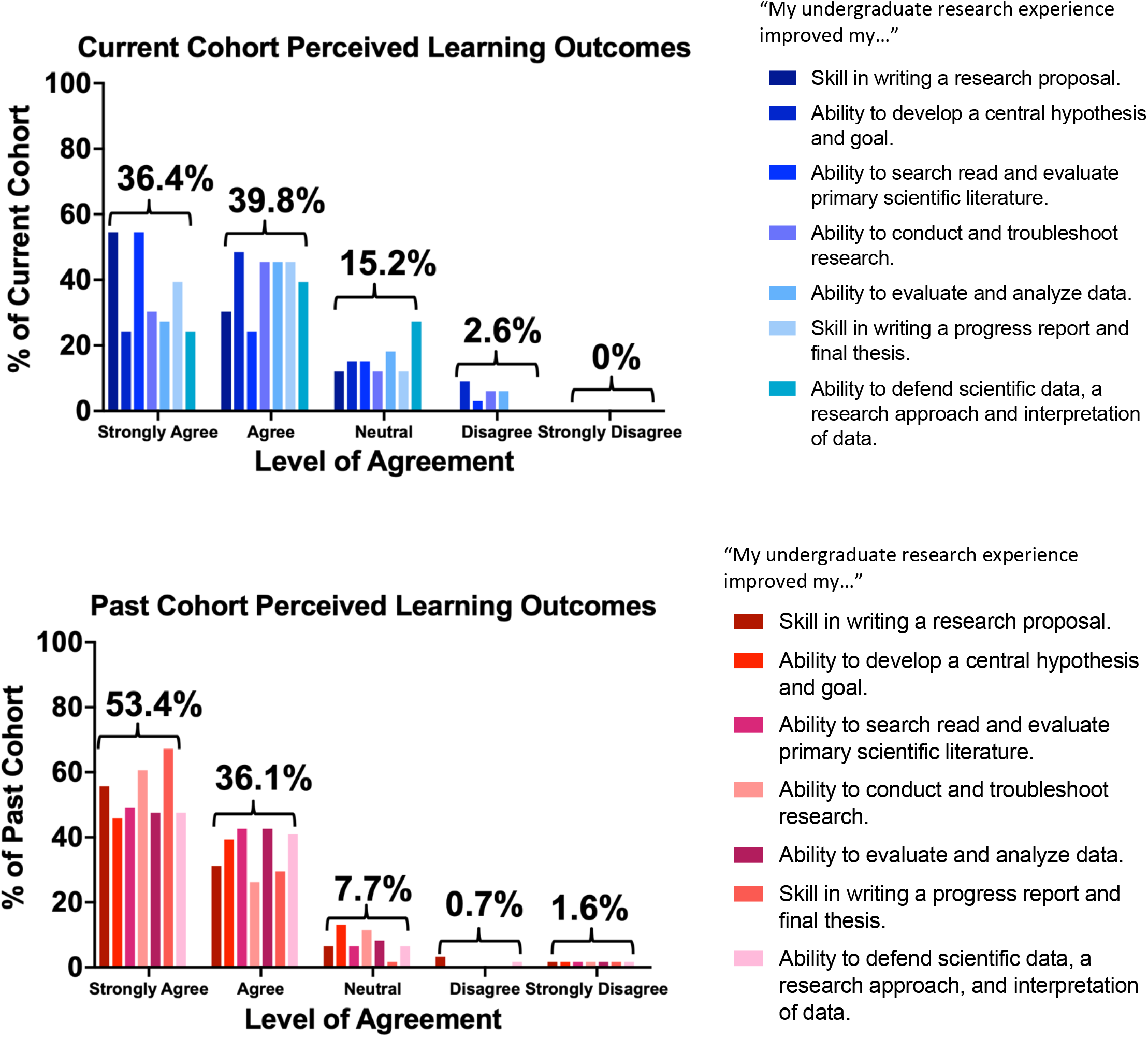
Measures of perceived learning outcomes divided by current cohort (2020/2021 academic year) and past cohort (5 academic years prior to 2020/2021)

### 4.4 Emergent Themes and Recommendations

Although thematic analysis has not yet been finalized due to time constraints, preliminary analysis has revealed recurrent points of complaint and feedback among both the current and past cohort (Tables 4 and 5). Themes that were identified by both the current and past cohort include: 1) positive experiences & gain of soft skills, 2) Bio4999 lecture content was not relevant or helpful, 3) lack of guidance & feelings of isolation, and 4) the need to clarify expectations to research supervisors.

**Table 4.**
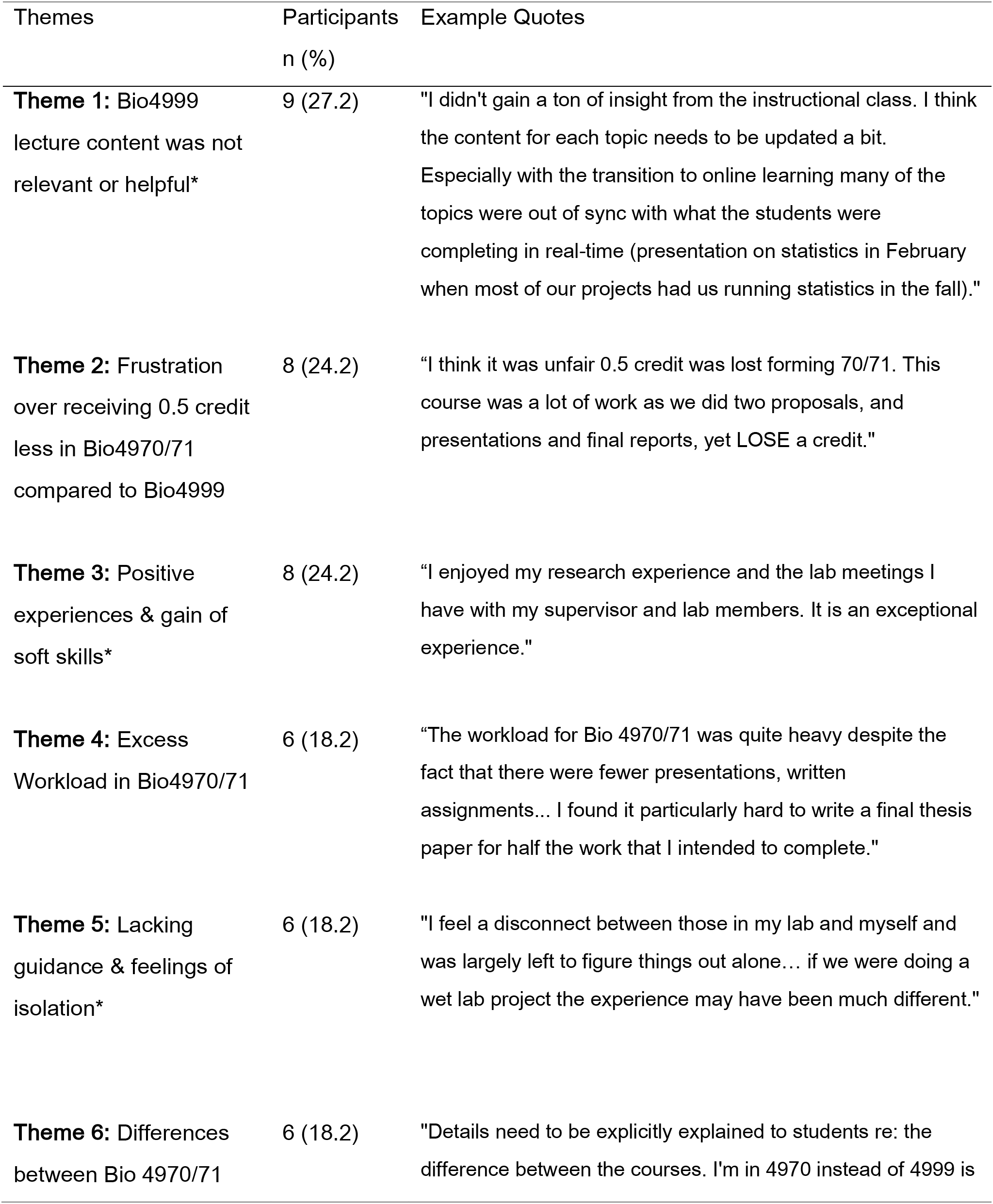
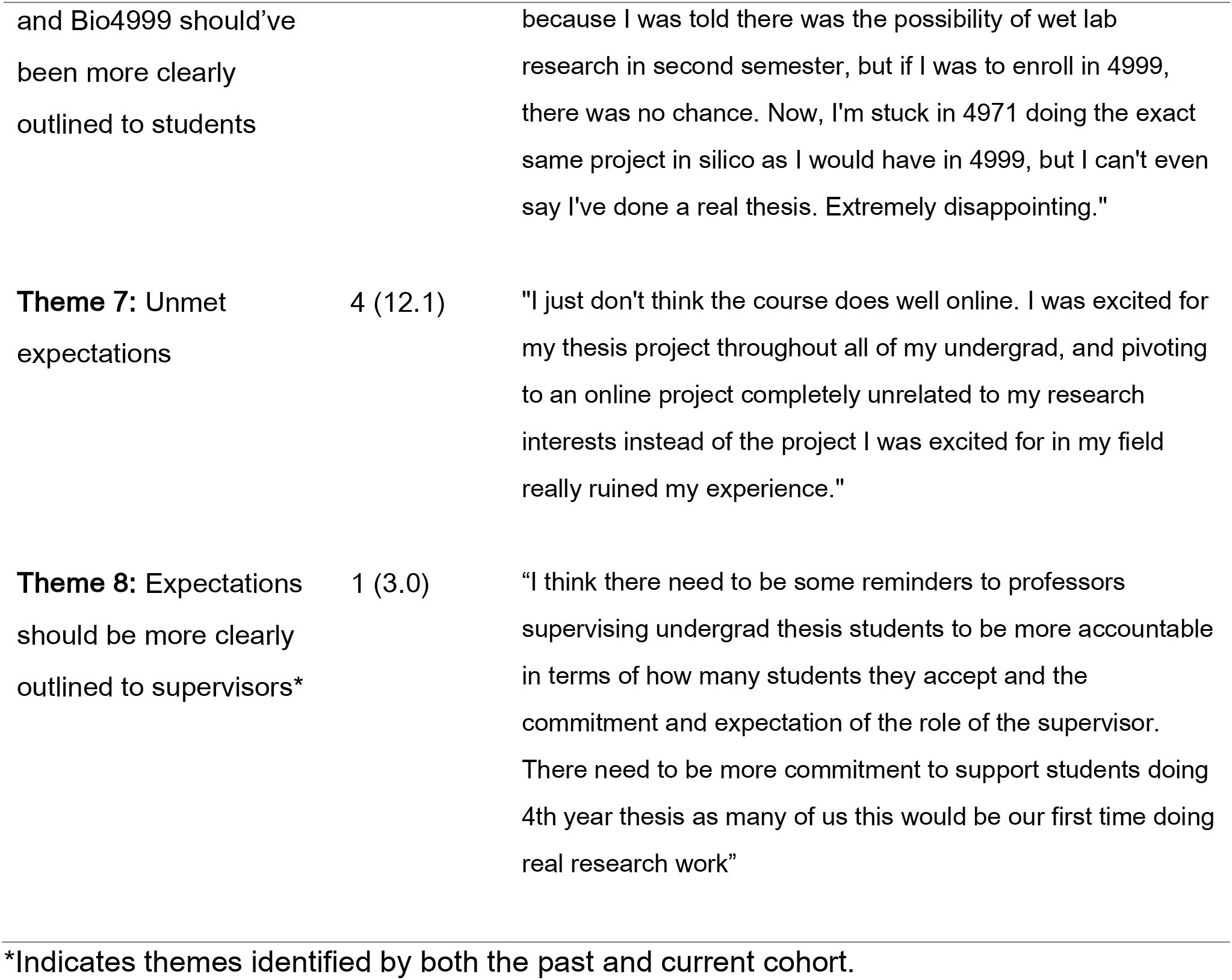
Preliminary themes from current cohort’s (2020/2021 academic year) survey responses.

| Themes | Participants<br>n (%) | Example Quotes |
| --- | --- | --- |
| **Theme 1:** Bio4999 lecture content was not relevant or helpful* | 9 (27.2) | "I didn't gain a ton of insight from the instructional class. I think the content for each topic needs to be updated a bit. Especially with the transition to online learning many of the topics were out of sync with what the students were completing in real-time (presentation on statistics in February when most of our projects had us running statistics in the fall)." |
| **Theme 2:** Frustration over receiving 0.5 credit less in Bio4970/71 compared to Bio4999 | 8 (24.2) | "I think it was unfair 0.5 credit was lost forming 70/71. This course was a lot of work as we did two proposals, and presentations and final reports, yet LOSE a credit." |
| **Theme 3:** Positive experiences & gain of soft skills* | 8 (24.2) | "I enjoyed my research experience and the lab meetings I have with my supervisor and lab members. It is an exceptional experience." |
| **Theme 4:** Excess Workload in Bio4970/71 | 6 (18.2) | "The workload for Bio 4970/71 was quite heavy despite the fact that there were fewer presentations, written assignments... I found it particularly hard to write a final thesis paper for half the work that I intended to complete." |
| **Theme 5:** Lacking guidance & feelings of isolation* | 6 (18.2) | "I feel a disconnect between those in my lab and myself and was largely left to figure things out alone... if we were doing a wet lab project the experience may have been much different." |
| **Theme 6:** Differences between Bio 4970/71 | 6 (18.2) | "Details need to be explicitly explained to students re: the difference between the courses. I'm in 4970 instead of 4999 is |
| and Bio4999 should've<br>been more clearly<br>outlined to students |  | because I was told there was the possibility of wet lab<br>research in second semester, but if I was to enroll in 4999,<br>there was no chance. Now, I'm stuck in 4971 doing the exact<br>same project in silico as I would have in 4999, but I can't even<br>say I've done a real thesis. Extremely disappointing." |
| **Theme 7:** Unmet<br>expectations | 4 (12.1) | "I just don't think the course does well online. I was excited for<br>my thesis project throughout all of my undergrad, and pivoting<br>to an online project completely unrelated to my research<br>interests instead of the project I was excited for in my field<br>really ruined my experience." |
| **Theme 8:** Expectations<br>should be more clearly<br>outlined to supervisors* | 1 (3.0) | "I think there need to be some reminders to professors<br>supervising undergrad thesis students to be more accountable<br>in terms of how many students they accept and the<br>commitment and expectation of the role of the supervisor.<br>There need to be more commitment to support students doing<br>4th year thesis as many of us this would be our first time doing<br>real research work" |

\*Indicates themes identified by both the past and current cohort.

**Table 5.**
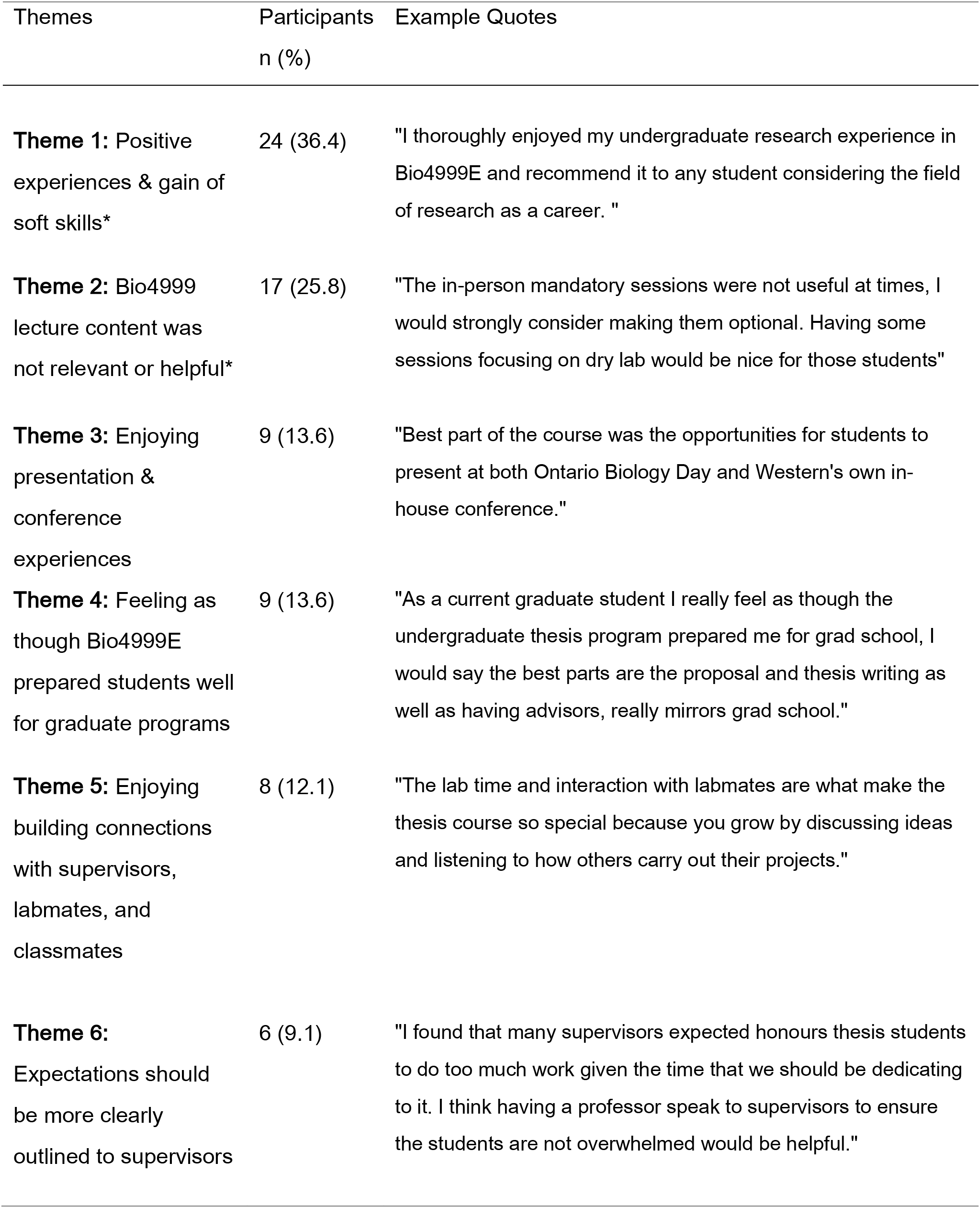
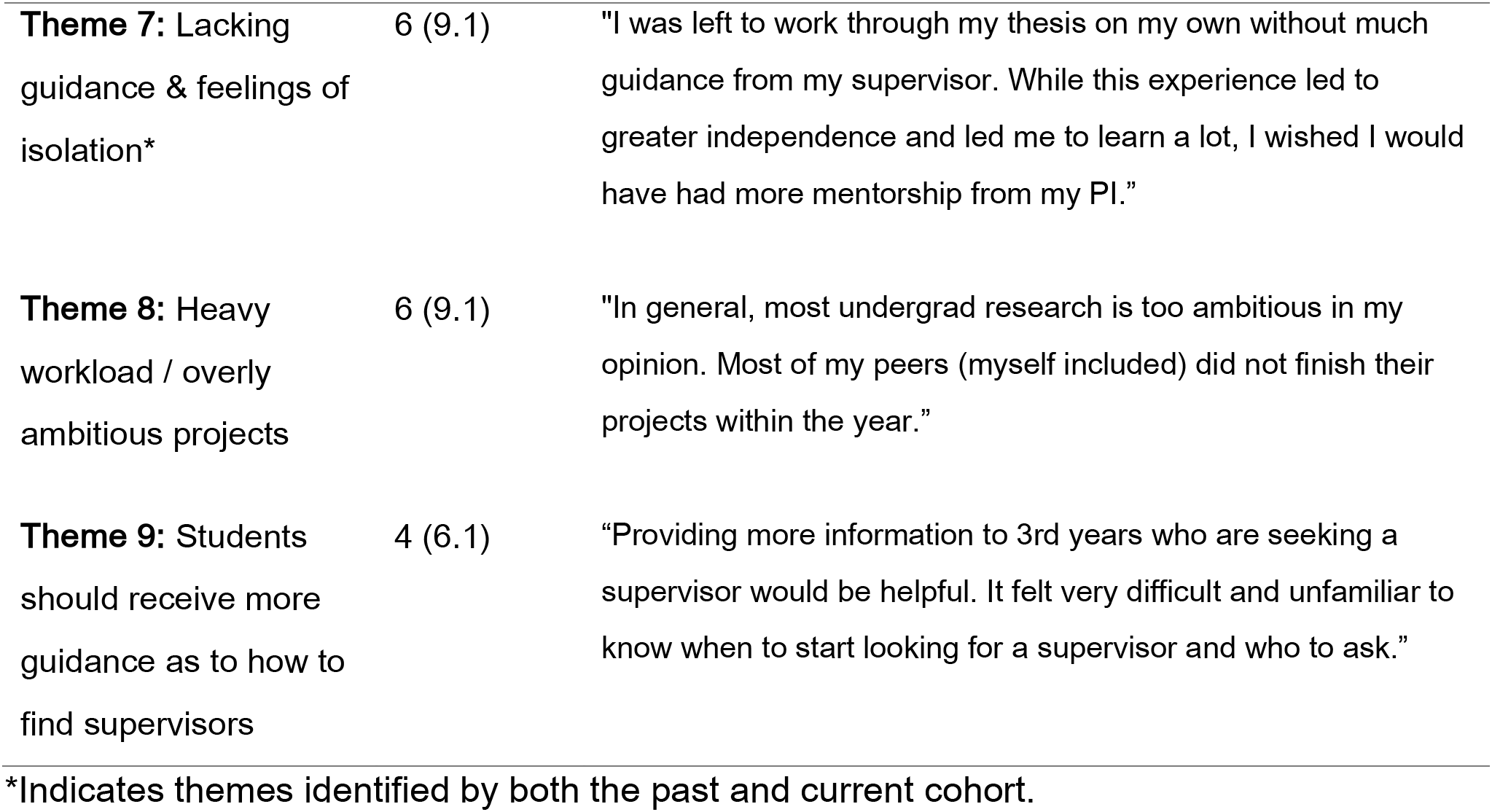
Preliminary themes from past cohort’s (5 academic years prior to 2020/2021) survey responses.

| Themes | Participants<br>n (%) | Example Quotes |
| --- | --- | --- |
| **Theme 1:** Positive experiences & gain of soft skills* | 24 (36.4) | "I thoroughly enjoyed my undergraduate research experience in Bio4999E and recommend it to any student considering the field of research as a career. " |
| **Theme 2:** Bio4999 lecture content was not relevant or helpful* | 17 (25.8) | "The in-person mandatory sessions were not useful at times, I would strongly consider making them optional. Having some sessions focusing on dry lab would be nice for those students" |
| **Theme 3:** Enjoying presentation & conference experiences | 9 (13.6) | "Best part of the course was the opportunities for students to present at both Ontario Biology Day and Western's own in-house conference." |
| **Theme 4:** Feeling as though Bio4999E prepared students well for graduate programs | 9 (13.6) | "As a current graduate student I really feel as though the undergraduate thesis program prepared me for grad school, I would say the best parts are the proposal and thesis writing as well as having advisors, really mirrors grad school." |
| **Theme 5:** Enjoying building connections with supervisors, labmates, and classmates | 8 (12.1) | "The lab time and interaction with labmates are what make the thesis course so special because you grow by discussing ideas and listening to how others carry out their projects." |
| **Theme 6:** Expectations should be more clearly outlined to supervisors | 6 (9.1) | "I found that many supervisors expected honours thesis students to do too much work given the time that we should be dedicating to it. I think having a professor speak to supervisors to ensure the students are not overwhelmed would be helpful." |
| **Theme 7:** Lacking guidance & feelings of isolation* | 6 (9.1) | "I was left to work through my thesis on my own without much guidance from my supervisor. While this experience led to greater independence and led me to learn a lot, I wished I would have had more mentorship from my PI." |
| **Theme 8:** Heavy workload / overly ambitious projects | 6 (9.1) | "In general, most undergrad research is too ambitious in my opinion. Most of my peers (myself included) did not finish their projects within the year." |
| **Theme 9:** Students should receive more guidance as to how to find supervisors | 4 (6.1) | "Providing more information to 3rd years who are seeking a supervisor would be helpful. It felt very difficult and unfamiliar to know when to start looking for a supervisor and who to ask." |

\*Indicates themes identified by both the past and current cohort.

### 4.5 Stakeholders

A total of 58 stakeholders were identified with 32 (55.2%) in academia, 9 (15.5%) in media, 8 (13.8%) in government, 5 in the public (8.6%), and 4 in industry (6.9%) (Figure 7). Furthermore, a total of 107 connections were made between key stakeholders.

**Figure 7.**
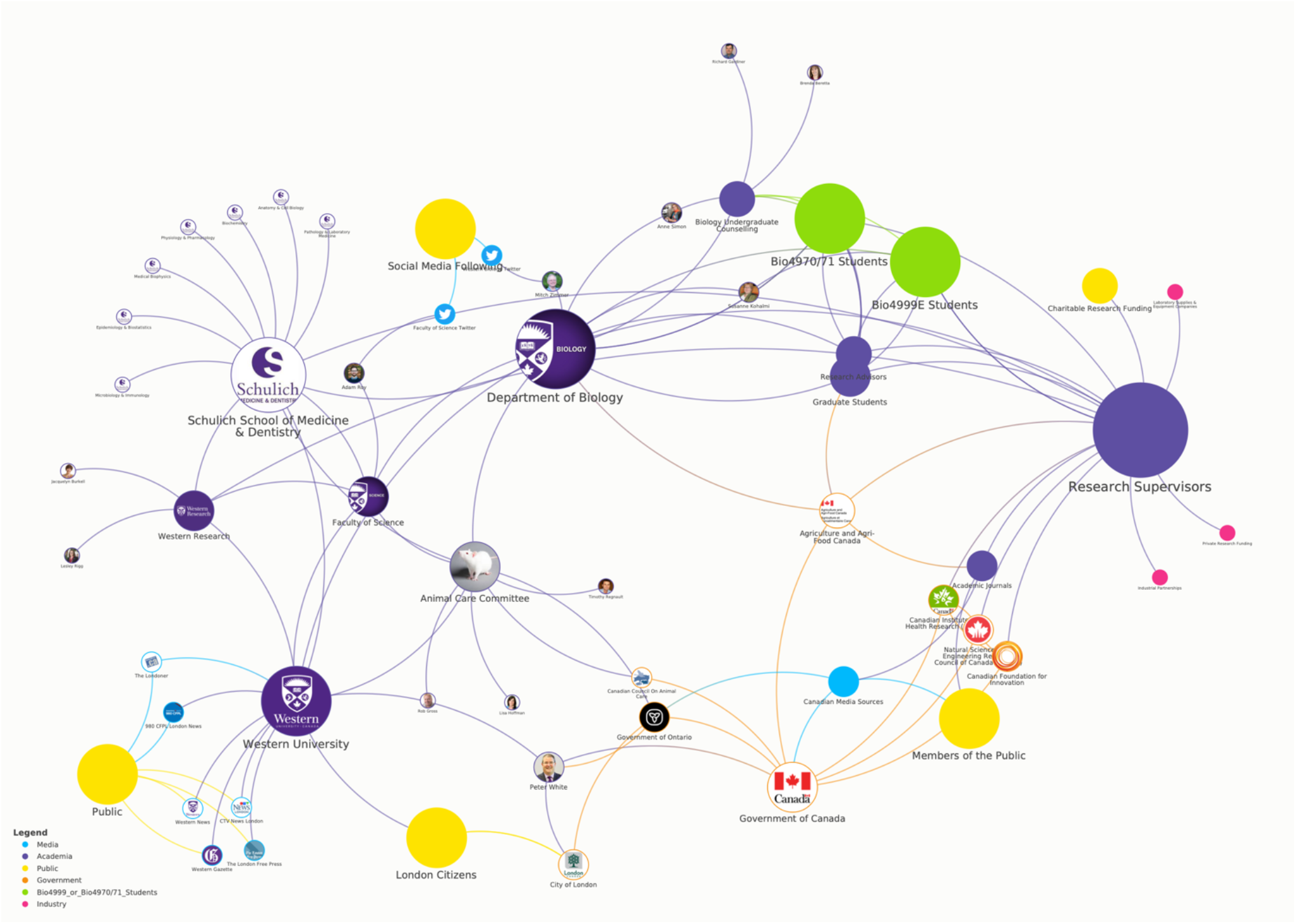
Key stakeholders related to undergraduate research in the Department of Biology at Western University. An interactive view is available at: https://embed.kumu.io/e1145410f796296d89f8aec78747d3aa

Key stakeholders within academia included research supervisors, Bio4999 and Bio4970/71 students, the Department of Biology, Schulich School of Medicine and Dentistry, Western University, the Animal Care Committee at Western, Western Research among others. The Animal Care committee is particularly relevant to this study as it serves as the governing body ensuring that all researchers at Western University adhere to the mandatory federal (Canadian Council on Animal Care) and provincial (Ontario Animals in Research Act) guidelines and standards for the use of animal models in research.

Key media stakeholders include the London News, CTV News London, the London Free Press, 980 CFPL London News, the Western Gazette, and Western News. Media often provided a direct connection to members of the public, thus promoting the spread of undergraduate research findings. An important note is that while only 5 stakeholders representing members of the public have been identified, each of these stakeholders constitutes a large group of people. For this reason, several of the bubbles representing members of the public on the stakeholder map have been increased in size to demonstrate the large number of individuals represented and the importance of the public in this particular topic.

Governmental stakeholders include the City of London, the Government of Ontario, and the Government of Canada, along with associated governmental agencies such as the Canadian Council on Animal Care, Agriculture and Agri-Food Canada, and the Natural Science and Engineering Council of Canada. Government at all three levels (local, regional, and national) interact with Western University on a regular basis and can have significant influence over the amount of funding received for instruction, research, student support programs, and more. Finally, three industry stakeholders were identified, and they include private institutions funding the work of researchers at Western University, industry members who form partnerships with researchers at Western in order to address specific environmental or human health concerns, and laboratory supplies and equipment companies that play a vital role in allowing research to function.

The 5 stakeholders with the most connections included: research supervisors (16 connections), the Department of Biology at Western University (14), Western University (13), Schulich School of Medicine and Dentistry (13), and the Government of Canada (8 connections). Notably, Bio4999 and Bio4970/71 students each had 6 direct connections, though were within 3 connections of a total of 37 stakeholders.

## 5. Discussion

### 5.1 Principal Findings

Despite COVID-19 related restrictions, the number of undergraduate research projects performed in the 2020/2021 academic year in the Department of Biology at Western University remained within range of enrollment in the 5 previous academic years (2015/2016, 2016/2017, 2017/2018, 2019/2020 and 2020/2021) (Figure 1). This indicates that the Department of Biology was able to accomplish its goal of accommodating undergraduate researchers despite reduced laboratory access on campus due to the COVID-19 pandemic.

Although expected, the shift toward dry research among the current cohort is quite striking (87.8% of the current cohort completed dry research, compared to 15.2% of the past cohort) (Figure 2.). Despite having the option to complete in-person research if students opted to take Bio4970/71, very few students ended up completing in-person research. This was likely due to fluctuating COVID-19 guidelines leading to varying degrees of restricted laboratory access at different points of the academic year. Interestingly, the number of projects using animal models was not dramatically affected by the shift to dry research (45.5% in the current cohort compared to 37.9% in the past; Figure 3), suggesting that COVID-19 likely promoted the use of data that had previously been collected from animal models, though had not yet been analyzed. From a One Health perspective, the shift towards dry research has repercussions for the type of knowledge generated and exchanged by researchers. Indeed, on a nation-wide or global scale, if COVID-19 restrictions remain in place for a prolonged period of time this could lead to severe repercussions to the types of studies conducted and the type of results generated. This could in turn impact our ability to effectively survey, detect, respond to, or treat various environmental, animal, or human health problems. Indeed, Weiner, Balasubramaniam, Shah, & Javier, 2020 argued that in the long-term, the COVID-19 pandemic will likely lead to reallocation of research dollars at the expense of research areas funded prior to the pandemic. This will require scientists, research institutions, patients, and other key stakeholders to engage in discussion and decisions regarding research goals and funding allocations. Moreover, Dall’Olio et al., 2020 point out that while COVID-19 might have led to a slight increase in research productivity in the short term as scientists had the opportunity to work on submitting long-standing analysis and conduct back-burned data analysis, it is likely that a lack of new data collection will lead to decreased productivity and research output in the future.

Compared to the past cohort, the proportion of students in the current cohort who had research supervisors outside of the Department of Biology decreased slightly (from 28.1% in the past cohort to 9.1% in the current cohort). While prior research has suggested that the COVID-19 pandemic has prompted increased collaboration between scientists in different fields (Korbel & Stegle, 2020; Radecki & Schonfeld, 2020; Weiner, Balasubramaniam, Shah, & Javier), this finding suggests that in relation to undergraduate research this might not be the case. Indeed, as resources are already spread thin and researchers are being forced to prioritize who has access to their laboratory space and/or research equipment, researchers may be less likely to extend resources to undergraduate students who are not already in their laboratory or in their field of research. Collaboration across sectors is vital to One Health, thus this finding raises concern (Centers for Disease Control and Prevention, 2019; World Health Organization, 2017).

The current cohort indicated overall lower levels of satisfaction and enjoyment with their undergraduate research experience, though perceived learning outcomes were fairly consistent with previous years (Figures 4 & 6). The finding of decreased levels of satisfaction and enjoyment among undergraduate students conducting research amidst the COVID-19 pandemic is in line with previous research (Deveau, Wang, & Small, 2020; Trego, Nadybal, Morales, Collins, & Grineski, 2020; Wang, Bauer, Burmeister, Hanauer, & Graham, 2020). To our knowledge, no other research has examined the impact of the COVID-19 pandemic on undergraduate researchers’ perception of having achieved learning outcomes.

Several factors may be contributing to the decreased levels of satisfaction among undergraduate students, including frustrations with online learning and research platforms, increased feelings of isolation due to reduced communication and interaction with research supervisors, labmates, course professors, and fellow undergraduate researchers, increased workload, decreased engagement and situational interests, and more (Adedoyin & Soykan, 2020; Christian, McCarty & Brown, 2020; Deveau, Wang, & Small, 2020; Gillis & Krull, 2020; Mukhtar et al., 2020; Trego, Nadybal, Morales, Collins, & Grineski, 2020; Wang, Bauer, Burmeister, Hanauer, & Graham, 2020). Indeed, several of these factors were directly identified by students in the free-ended portion of the survey (Table 4). An interesting finding was that students who indicated that their research projects had undergone major alterations reported lower levels of satisfaction compared to those who indicated minor or no alterations. This suggests a potential relationship between whether students’ expectations for their research project were met and overall satisfaction. Previous research has demonstrated that higher levels of enjoyment among students in online-learning environments are associated with increased motivation and engagement, and achievement of higher-level learning outcomes (Gomez, Wu, & Passerini, 2010). Thus, there is a need to incorporate creative online learning strategies targeting student enjoyment into online undergraduate research courses.

Preliminary analysis of qualitative data has revealed a wide range of concerns among students (Tables 4 and 5). Several of the student’s complaints were related to miscommunication or lack of communication (which subsequently led to feelings of isolation), thus demonstrating the importance of examining key stakeholder involvement and ensuring strong lines of communication and collaboration between stakeholders. For instance, students recommended increased communication between Bio4970/71 and Bio4999 professors (Dr. Anne Simon and Dr. Susanne Kohalmi, respectively) and research supervisors to ensure that students are receiving adequate support from their supervisors and are not being expected to complete an excessive amount of work. Moreover, while students in the past cohort indicated that having the opportunity to form connections with graduate students in their lab was one of the best features of their undergraduate research experience, students in the current cohort indicated a “disconnect” between themselves and the other members of their lab (Tables 4 and 5). Thus, providing opportunities for undergraduate students to directly interact and connect with graduate students despite being in an online environment may be of great value in future years. Overall, students provided invaluable feedback that can be used to develop specific recommendations to improve the quality of undergraduate research in the Department of Biology in the future.

Finally, a total of 58 stakeholders and 107 connections were identified, demonstrating the collaborative nature of undergraduate research and the wide context within which undergraduate research takes place in (Figure 7). Indeed, while undergraduate research can have the appearance of being isolated within academic communities, the stakeholder map developed for this study demonstrates that this is not the case. Rather, undergraduate research involves stakeholders across academia, government, media, industry, and the public. Moreover, the stakeholder map developed for this study demonstrates that there are several ways in which study findings produced by an undergraduate researcher can reach a governmental official, industry representative, media representative, or member of the public, thus highlighting the impact and value of undergraduate research.

### 5.2 Limitations

Due to the small sample size (n=99), findings may have limited generalizability. However, we are confident that findings from this study are highly relevant and useful to the Department of Biology at Western University, and we believe there are still valuable lessons from this study that that could be applied to other departments and universities. Survey results may be subject to non-response and recall bias. Individuals who were extremely dissatisfied or satisfied with their undergraduate research experience may be more likely to respond to the survey compared to those with more neutral feelings about their experience. Furthermore, individuals who completed their undergraduate research project in the 4 academic years prior to the 2020/2021 academic year may be more likely to reflect positively on their experience now that they are in graduate or professional school or have begun their professional careers. Finally, the negative impact of the COVID-19 pandemic on students overall mental health and wellbeing might have led to an increased likelihood to reflect and report negatively on undergraduate research experience.

### 5.3 Strengths and Significance

This study provided a timely assessment of the impact of the COVID-19 pandemic on undergraduate researchers. As a great deal of uncertainty remains around COVID-19, it is foreseeable that online or blended learning might remain the norm for years to come. The insights gained by surveying students will help improve the quality of undergraduate research in the Department of Biology and will inform decisions about what elements of online learning to maintain moving forward. The results of this study may also be of use to other departments at Western University and other educational institutions who are looking to improve their own undergraduate research courses or incorporate experiential-based online learning strategies into existing courses. Undergraduate research has numerous benefits for both students and society at large, including allowing students to gain clarity about their career path, providing the university with publications and conference presentations, increasing the visibility of the scientific community, and more. Therefore, it is important that courses offering undergraduate research opportunities are of high quality.

To our knowledge, this is the only study that has examined the impact of COVID-19 on undergraduate research, or even research in general, through the lens of One Health. Using the One Health approach for this study allowed us to situate our study findings within a wider context, considering the impact of COVID-19 on post-secondary students’ learning and social environments and on research productivity in both the short and long-term. Ultimately, the use of the One Health approach led to a deeper consideration of the potential human, animal, and environmental health repercussions associated with COVID-19’s impact on undergraduate research.

### 5.4 Future Directions

Future work should focus on the development of a list of specific recommendations to improve the quality of undergraduate research in the future and to help universities, faculty members, and students prepare for further disruptions to undergraduate research as a result of the COVID-19 pandemic or other causes. As well, studies quantifying the long-term impacts of the COVID-19 pandemic on research in general and on undergraduate research specifically are needed. These studies should assess impacts of the COVID-19 pandemic on the amount of research performed, the type of research performed, the levels of satisfaction and enjoyment of researchers, and the degree of collaboration between different sectors.

## 6. Acknowledgments

I would like to thank my wonderful supervisor, Dr. Anne Simon, as well as all the other members of the Simon lab for their guidance and support throughout this research project. In particular I would like to thank Wes Robinson and Ryley Yost, the graduate students, for their help. Thank you to Dr. Olea Popelka, my fellow One Health peers, and the Department of Pathology for supplying me with the tools and support necessary to complete my research. Finally, I would like to thank my advisors Dr. Susanne Kohalmi for her time and expertise.

## 7. Author Contributions

A.C. designed study and collected data, developed figures and tables, wrote the first draft of the paper. A.S. conceptualized the project, supervised and contributed directly to the design and collection of data. S.K. contributed to study design.

## Appendix 1. Current Cohort Survey

### LETTER OF INFORMATION AND CONSENT

Impact of the Pandemic on Undergraduate Research in Biology

#### Researchers

Anne F. Simon, PhD

Principal Investigator

Associate Chair (Undergraduate), Department of Biology, Faculty of Science Western University, 519-661-2111 x80084

#### Ava Chaplin

Co-Investigator

One Health Honor’s Thesis Student, Department of Pathology & Laboratory Medicine Western University, phone number: 519-831-9884

#### Contact Information

Please contact Ava Chaplin by email or telephone (519-831-9884) with any questions you might have.

#### Invitation to Participate

You are being asked to participate in a study regarding the impact of the COVID-19 pandemic on the experience of students completing an undergraduate research project in the Department of Biology. The purpose of this letter is to provide you with information required for you to make an informed decision regarding participation in this research.

#### Study Purpose

This study is being done to investigate the effect of the reduced access to research laboratories as a consequence of the COVID-19 pandemic on the experience of students conducting an independent research project for the Bio4999 or Bio4970/71 courses.

#### Inclusion Criteria

Individuals who are currently enrolled in Bio4999E or Bio4971G at Western University, or who completed Bio4999 in the past 4 years are eligible to participate in this study.

#### Procedures

If you agree to participate, after reading this letter, you will be asked to fill out a ∼25 items survey about your experience as a student researcher that should take approximately 10-15 minutes to complete. It is not mandatory to answer every question, and you can skip any question that you choose.

#### Risks & Benefits to Participants

You will be asked to reflect on your experience as a student researcher, which may trigger minor feelings of stress or anxiety, or alternatively make your see your experience in a positive way. There is no compensation for completing the survey.

#### Benefits to Society

Your responses will provide us with valuable information as to how we can improve future undergraduate research opportunities in the Department of Biology.

#### Confidentiality

All responses to this survey are anonymous. Personal identifiers (e.g. name, address, date of birth, etc.) will NOT be collected. Anonymous data files will stored on a password protected computer, and will only be accessed by the principal investigator and co-investigator. All data will be retained for seven years as per regulatory guidelines. Your survey responses will be collected anonymously through a secure online survey platform called Qualtrics. Qualtrics uses encryption technology and restricted access authorizations to protect all data collected. In addition, Western’s Qualtrics server is in Ireland, where privacy standards are maintained under the European Union safe harbour framework. The data will then be exported from Qualtrics and securely stored on Western University’s server.

#### Rights as a Participant

Your decision to participate in this study is completely voluntary and will have no impact on your academic standing. You do not waive any legal rights by consenting to this survey.

You may refuse to participate and refuse to answer any questions at any point in this study. If at any point you decided to withdraw your participation while completing the survey you may do so simply by closing your browser window. Once data has been submitted, you will be unable to withdraw from the study.

If you have any questions about your rights as a research participant or the conduct of this study, you may contact The Office of Human Research Ethics (519) 661-3036. This office oversees the ethical conduct of research at Western University and is not a part of the research team.

QI Consent

○ This study has been explained to me and any questions I had have been answered. I know that I may leave the survey at any time. I understand that by responding to the survey questions I am indicating voluntary agreement to participate. (1)

Q1 Please indicate which of the following courses you are currently enrolled in:

○ I am currently enrolled in Bio4999E (1)
○ I am currently enrolled in Bio4971G (2)
○ Prefer not to answer (3)

Q2 What Honors Specialization do you plan on graduating with?

________________________________

Q3 Prior to June 2020, were you planning on living in London for the 2020/2021 Academic School Year?

○ Yes (1)
○ No (2)
○ Maybe (3)
○ Prefer not to answer (4)

Q4 Are you living in London for the 2020/2021 Academic School Year?

○ Yes, full time (1)
○ Yes, part time (2)
○ No (3)
○ Prefer not to answer (4)

Q5 Prior to Bio4999 or Bio4970/71 did you have any research experience?

○ Yes (1)
○ No (2)
○ Prefer not to answer (3)

Q6 If you answered ‘Yes’ to the question above, prior to Bio4999 or Bio4970/71 had you shared any research results? Select all that apply.

○ Yes, I presented an oral poster. (1)
○ Yes, I was made (co-)author on a peer-reviewed publication. (2)
○ No, I have not shared any research results. (3)
○ Prefer not to answer (4)

Q7 Before the start of the project in September, when did you find your supervisor?

○ May-August 2020 (1)
○ January-April 2020 (2)
○ September-December 2019 (3)
○ May-August 2019 (4)
○ Before May 2019 (5)
○ Prefer not to answer (6)

Q8 What department is your research supervisor in?

________________________________

Q9 Did your research project undergo any changes between registration in Bio4999 or Bio4970 in June 2020 and the first proposal deadline in October 2020?

○ Yes, my project underwent major alterations, for example a shift in research focus or a switch from wet to dry research. (1)
○ Yes, my project underwent minor alterations, for example an altered hypothesis or updated project goals. (2)
○ No, my project had no alterations. (3)
○ Prefer not to answer (4)

Q10 When you first met with your supervisor and discussed plans for your thesis project, did you intend to complete a **wet** (e.g. involving experimental work taking place in a laboratory), **dry** (e.g. *in silico*, bioinformatics research, data analysis, or bibliography research completed on a computer) or **field-based** (e.g. involving collection of primary data from a natural environment) research project?

○ Wet (1)
○ Dry (2)
○ Field (3)
○ More than one of the above. Please state the combination: (4)
________________________________
○ Prefer not to answer (5)

Q11 What type of research did you end up completing?

○ Wet (1)
○ Dry (2)
○ Field (3)
○ More than one of the above. Please state the combination: (4)
________________________________
○ Prefer not to answer (5)

Q12 At the end of Bio4999 or Bio4970/71, do you anticipate that your results will be shared?

○ Yes, I will present a poster or an abstract at a conference other than Ontario Biology Day. (1)
○ Yes, my results will be presented by labmates at a conference. (2)
○ Yes, I will be an author on a peer-reviewed publication. (3)
○ No (4)
○ Prefer not to answer (5)

Q13 Does your research project involve the use of an animal model?

○ Yes (1)
○ No (2)
○ Prefer not to answer (3)

Q14 If you answered ‘Yes’ to the question above, what type of animal model are you using?

________________________________

Q15 **Please indicate how strongly you agree or disagree with the following statements regarding your experience in Bio4999 and/or Bio4970/71.** If for any reason you prefer not to answer, or consider the question irrelevant to you, please choose the “Not applicable / Prefer not to answer” option.

|  | Strongly<br>Disagree<br>(1) | Disagree<br>(2) | Neutral<br>(3) | Agree<br>(4) | Strongly<br>Agree (5) | N/A or<br>Prefer not<br>to answer<br>(6) |
| --- | --- | --- | --- | --- | --- | --- |
| I am satisfied with my research experience. (1) | ☐ | ☐ | ☐ | ☐ | ☐ | ☐ |
| My research experience met my expectations. (2) | ☐ | ☐ | ☐ | ☐ | ☐ | ☐ |
| My research experience was enjoyable. (3) | ☐ | ☐ | ☐ | ☐ | ☐ | ☐ |
| This experience was a good way of learning about the process of scientific research. (4) | ☐ | ☐ | ☐ | ☐ | ☐ | ☐ |
| This experience had a positive effect on my interest in science. (5) | ☐ | ☐ | ☐ | ☐ | ☐ | ☐ |
| I would recommend Bio4970/71 or Bio4999 to another undergraduate student in Biology. (6) | ☐ | ☐ | ☐ | ☐ | ☐ | ☐ |

Q16 In this section of the survey, you will be asked to consider a variety of possible benefits from your research experience. **Please indicate how strongly you agree or disagree with the following statements. Please assume that each statement begins with “As a result of my experience in Bio4999 or Bio4970/71, I improved my…”.** If for any reason you prefer not to answer, or consider the question irrelevant to you, please choose the “Not applicable / Prefer not to answer” option.

|  | Strongly<br>Disagree<br>(1) | Disagree<br>(2) | Neutral (3) | Agree (4) | Strongly<br>Agree (5) | N/A or<br>Prefer not<br>to answer<br>(6) |
| --- | --- | --- | --- | --- | --- | --- |
| Skill in writing a research proposal. (1) |  | ☐ | ☐ | ☐ | ☐ | ☐ |
| Ability to develop a central hypothesis and goal. (2) |  | ☐ | ☐ | ☐ | ☐ | ☐ |
| Ability to search, read, and evaluate primary scientific literature. (3) |  | ☐ | ☐ | ☐ | ☐ | ☐ |
| Ability to conduct and troubleshoot research. (4) |  | ☐ | ☐ | ☐ | ☐ | ☐ |
| Ability to evaluate and analyze data. (5) |  | ☐ | ☐ | ☐ | ☐ | ☐ |
| Skill in writing a progress report and final thesis. (6) |  | ☐ | ☐ | ☐ | ☐ | ☐ |
| Ability to defend scientific data, a research approach, and interpretation of data. (7) |  | ☐ | ☐ | ☐ | ☐ | ☐ |

Q17 Do you have any constructive feedback for Bio4999E or Bio4970/71? For instance, in your experience what was the best part of the course and what could be changed?

________________________________
________________________________
________________________________
________________________________
________________________________

Q18 Please write any additional comments you have about your research experience here. Also, if you have any comments regarding the survey itself, please enter them here.

________________________________
________________________________
________________________________
________________________________
________________________________

## References

Adedoyin, O. B., & Soykan, E. (2020). Covid-19 pandemic and online learning: the challenges and opportunities. Interactive Learning Environments, 1–13. 10.1080/10494820.2020.1813180

Aucejo, E. M., French, J., Araya, M. P. U., & Zafar, B. (2020). The Impact of COVID-19 on Student Experiences and Expectations: Evidence from a Survey. Journal of Public Economics, 191(104271), 104271. 10.1016/j.jpubeco.2020.104271

Cao, W., Fang, Z., Hou, G., Han, M., Xu, X., Dong, J., & Zheng, J. (2020). The psychological impact of the COVID-19 epidemic on college students in China. Psychiatry Research, 287(112934),. 10.1016/j.psychres.2020.112934

Centers for Disease Control and Prevention. (2019). One Health. Retrieved from Centers for Disease Control and Prevention website: https://www.cdc.gov/onehealth/index.html

Christian, D. D., McCarty, D. L., & Brown, C. L. (2020). Experiential Education during the COVID-19 Pandemic: A Reflective Process. Journal of Constructivist Psychology, 33, 1–14. 10.1080/10720537.2020.1813666

Clarke, V., & Braun, V. (2016). Thematic analysis. The Journal of Positive Psychology, 12(3), 297–298. 10.1080/17439760.2016.1262613

Dall’Olio, R., Blacquiere, T., Bouga, M., Brodschneider, R., Carreck, N. L., Chantawannakul, P., … Neumann, P. (2020). COLOSS survey: global impact of COVID-19 on bee research. Journal of Apicultural Research, 59(5), 731–734. 10.1080/00218839.2020.1799646

Deveau, A. M., Wang, Y., & Small, D. J. (2020). Reflections on Course-Based Undergraduate Research in Organic and Biochemistry during COVID-19. Journal of Chemical Education, 97(9), 3463–3469. 10.1021/acs.jchemed.0c00787

Dumas, T. M., Ellis, W., & Litt, D. M. (2020). What Does Adolescent Substance Use Look Like During the COVID-19 Pandemic? Examining Changes in Frequency, Social Contexts, and Pandemic-Related Predictors. Journal of Adolescent Health. 10.1016/j.jadohealth.2020.06.018

Elmer, T., Mepham, K., & Stadtfeld, C. (2020). Students under lockdown: Comparisons of students’ social networks and mental health before and during the COVID-19 crisis in Switzerland. PLOS ONE, 15(7), e0236337. 10.1371/journal.pone.0236337

Gibbs, E. P. J. (2014). The evolution of One Health: a decade of progress and challenges for the future. Veterinary Record, 174(4), 85–91. 10.1136/vr.g143

Gillis, A., & Krull, L. M. (2020). COVID-19 Remote Learning Transition in Spring 2020: Class Structures, Student Perceptions, and Inequality in College Courses. Teaching Sociology, 48(4), 283–299. 10.1177/0092055x20954263

Gomez, E. A., Wu, D., & Passerini, K. (2010). Computer-supported team-based learning: The impact of motivation, enjoyment and team contributions on learning outcomes. Computers & Education, 55(1), 378–390. 10.1016/j.compedu.2010.02.003

Government of Canada. (2020). Guidance for post-secondary institutions during the COVID-19 pandemic. In. Retrieved from https://www.canada.ca/en/public-health/services/diseases/2019-novel-coronavirus-infection/guidance-documents/covid-19-guidance-post-secondary-institutions-during-pandemic.html

Kaparounaki, C. K., Patsali, M. E., Mousa, D.-P. V., Papadopoulou, E. V. K., Papadopoulou, K. K. K., & Fountoulakis, K. N. (2020). University students’ mental health amidst the COVID-19 quarantine in Greece. Psychiatry Research, 290, 113111. 10.1016/j.psychres.2020.113111

Kleiman, E. M., Yeager, A. L., Grove, J. L., Kellerman, J. K., & Kim, J. S. (2020). The real-time mental health impact of the COVID-19 pandemic on college students: An ecological momentary assessment study (Preprint). JMIR Mental Health, 7(12). 10.2196/24815

Korbel, J. O., & Stegle, O. (2020). Effects of the COVID-19 pandemic on life scientists. Genome Biology, 21(1). 10.1186/s13059-020-02031-1

Lerner, H., & Berg, C. (2015). The concept of health in One Health and some practical implications for research and education: what is One Health? Infection Ecology & Epidemiology, 5(1), 25300. 10.3402/iee.v5.25300

Linn, M. C., Palmer, E., Baranger, A., Gerard, E., & Stone, E. (2015). Undergraduate research experiences: Impacts and opportunities. Science, 347(6222), 1261757. 10.1126/science.1261757

Lopatto, D. (2007). Undergraduate Research Experiences Support Science Career Decisions and Active Learning. CBE—Life Sciences Education, 6(4), 297–306. 10.1187/cbe.07-06-0039

Lopatto, D., Tobias, S., Council On Undergraduate Research (U.S, & Research Corporation For Science Advancement. (2010). Science in solution : the impact of undergraduate research on student learning. Washington, D.C.: Council On Undergraduate Research; Tucson, Ariz.

Meda, N., Pardini, S., Slongo, I., Bodini, L., Zordan, M. A., Rigobello, P., … Novara, C. (2021). Students’ mental health problems before, during, and after COVID-19 lockdown in Italy. Journal of Psychiatric Research, 134, 69–77. 10.1016/j.jpsychires.2020.12.045

Mukhtar, K., Javed, K., Arooj, M., & Sethi, A. (2020). Advantages, Limitations and Recommendations for online learning during COVID-19 pandemic era. Pakistan Journal of Medical Sciences, 36(COVID19-S4). 10.12669/pjms.36.covid19-s4.2785

Murray, J. L. (2019). Undergraduate research for student engagement and learning. New York, Ny: Routledge.

Qiang, Z., Obando, A. G., Chen, Y., & Ye, C. (2020). Revisiting Distance Learning Resources for Undergraduate Research and Lab Activities during COVID-19 Pandemic. Journal of Chemical Education, 97(9), 3446–3449. 10.1021/acs.jchemed.0c00609

Radecki, J., & Schonfeld, R. (2020). The Impacts of COVID-19 on the Research Enterprise. 10.18665/sr.314247

Rea, L., & Parker, R. (2014). Designing and conducting survey research: A comprehensive guide. (4th ed.). Jossey-Bass.

Russell, S. H., Hancock, M. P., & McCullough, J. (2007). THE PIPELINE: Benefits of Undergraduate Research Experiences. Science, 316(5824), 548–549. 10.1126/science.1140384

Seymour, E., Hunter, A.-B., Laursen, S. L., & DeAntoni, T. (2004). Establishing the benefits of research experiences for undergraduates in the sciences: First findings from a three-year study. Science Education, 88(4), 493–534. 10.1002/sce.10131

Son, C., Hegde, S., Smith, A., Wang, X., & Sasangohar, F. (2020). Effects of COVID-19 on College Students’ Mental Health in the United States: Interview Survey Study. Journal of Medical Internet Research, 22(9),. 10.2196/21279

Speer, J. E., Lyon, M., & Johnson, J. (2021). Gains and Losses in Virtual Mentorship: A Descriptive Case Study of Undergraduate Mentees and Graduate Mentors in STEM Research during the COVID-19 Pandemic. CBE—Life Sciences Education, 20(2), ar14. 10.1187/cbe.20-06-0128

Totten, V. Y., Panacek, E. A., & Price, D. (1999). Basics of research (part 14) survey research methodology: Designing the survey instrument. Air Medical Journal, 18(1), 26–34. 10.1016/s1067-991x(99)90006-8

Trego, S., Nadybal, S., Morales, D., Collins, T., & Grineski, S. (2020). Initial Impacts of COVID-19 on Undergraduate Research (Undergraduate Dissertation). Retrieved from https://our.utah.edu/summer-symposium/trego/

Wang, C., Bauer, M., Burmeister, A. R., Hanauer, D. I., & Graham, M. J. (2020). College Student Meaning Making and Interest Maintenance During COVID-19: From Course-Based Undergraduate Research Experiences (CUREs) to Science Learning Being Off-Campus and Online. Frontiers in Education, 5. 10.3389/feduc.2020.590738

Wang, X., Hegde, S., Son, C., Keller, B., Smith, A., & Sasangohar, F. (2020). Investigating College Students’ Mental Health During the COVID-19 Pandemic: An Online Survey Study (Preprint). Journal of Medical Internet Research, 22(9). 10.2196/22817

Weiner, D. L., Balasubramaniam, V., Shah, S. I., & Javier, J. R. (2020). COVID-19 impact on research, lessons learned from COVID-19 research, implications for pediatric research. Pediatric Research, 88(2), 148–150. 10.1038/s41390-020-1006-3

World Health Organization. (2017, September 21). One Health. https://www.who.int/news-room/q-a-detail/one-health

